# A cationic liposome-formulated Toll Like Receptor (TLR)7/8 agonist enhances the efficacy of a vaccine against fentanyl toxicity

**DOI:** 10.1101/2025.01.08.631964

**Authors:** Fatima A Hamid, Nguyet-Minh Nguyen Le, Daihyun Song, Hardik Amin, Linda Hicks, Sophia Bird, Karthik Siram, Brooke Hoppe, Borries Demeler, Jay T. Evans, David Burkhart, Marco Pravetoni

## Abstract

The U.S. opioid epidemic is an extraordinary public health crisis that started in 1990 and significantly accelerated in the last decade. Since 2020, over 100,000 fatal drug overdoses have been reported annually, and 75% of those involved fentanyl and its analogs (F/FA). Accelerating the translation of innovative, effective, and safe treatments is needed to augment existing measures to counteract such a crisis. Active immunization against F/FA and other opioids represents a promising therapeutic and prophylactic strategy for opioid use disorder (OUD) and opioid-induced overdose toxicity. Previously we demonstrated that the anti-fentanyl vaccine comprising a fentanyl-based hapten (F) conjugated to the diphtheria cross-reactive material (CRM), admixed with the novel lipidated toll-like receptor 7/8 (TLR7/8) agonist INI-4001 adsorbed on Alhydrogel^®^ (alum) induced high-affinity fentanyl-specific polyclonal antibodies that protected against fentanyl-induced pharmacological effects in mice, rats, and mini-pigs. Here, INI-4001 was formulated into liposomes with different surface charges, and their impact on F-CRM adsorption, INI-4001 adjuvanticity, and vaccine efficacy were explored. Additionally, as the role of innate immunity in mediating the efficacy of addiction vaccines is largely unknown, we tested these formulations on the activation of innate immunity *in vitro*. Cationic INI-4001 liposomes surpassed other liposomal and aluminum-based formulations of INI-4001 by enhancing the efficacy of fentanyl vaccines and protecting rats against bradycardia and respiratory depression by blocking the distribution of fentanyl to the brain. Fentanyl vaccines adjuvanted with either cationic INI-4001 liposomes or the aqueous INI-4001 adsorbed to alum induced significant surface expression of co-stimulatory molecules and maturation markers in a murine dendritic cell line (JAWS II), while the former was superior in enhancing the macrophages surface expression of CD40, CD86 and inducible nitric oxide synthase (iNOS), indicative of maturation and activation. These results warrant further investigation of liposome-based formulations of TLR7/8 agonists for improving the efficacy of vaccines targeting F/FA and other opioids of public health interest.

**Graphical abstract:** 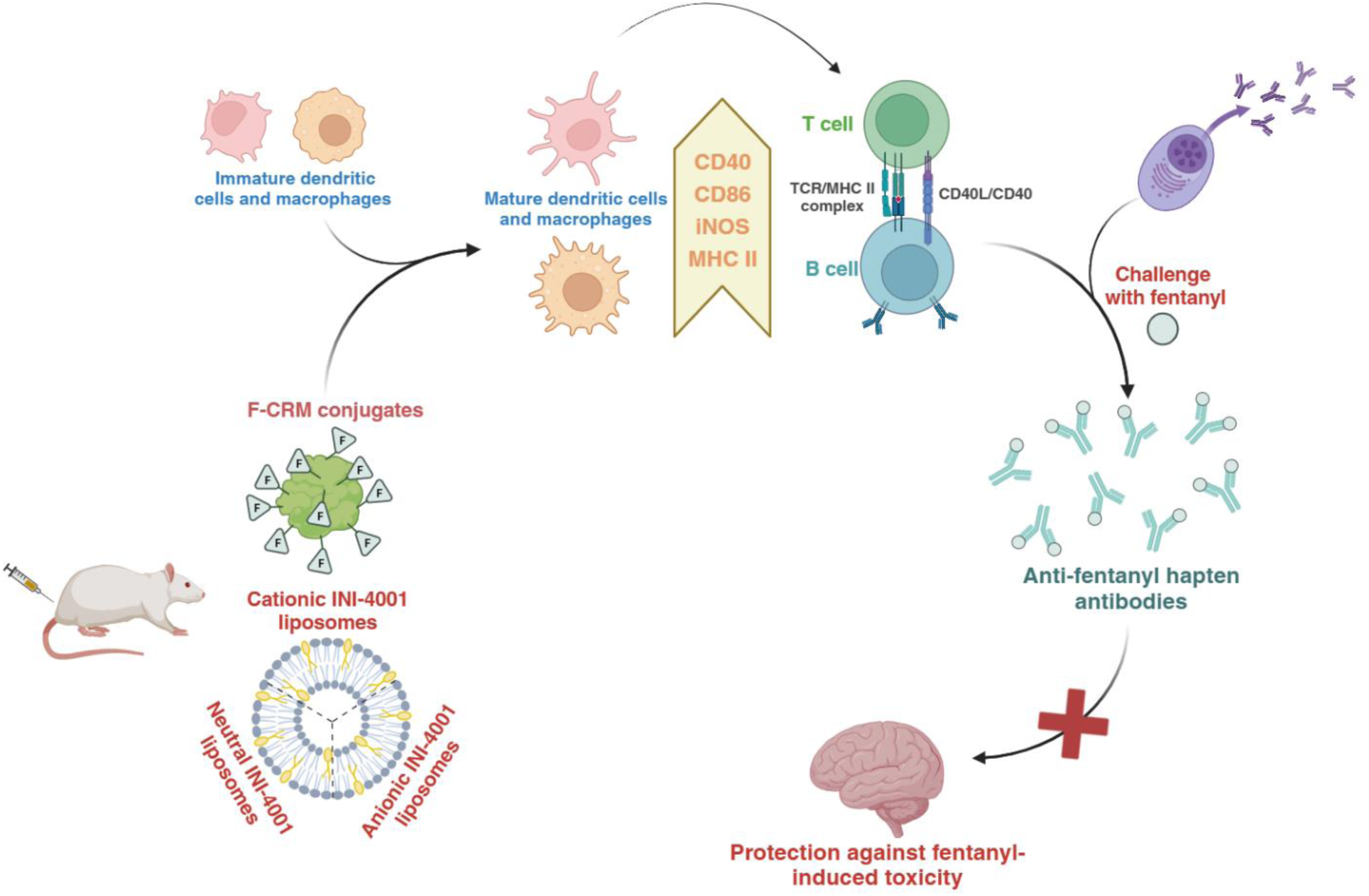

## 1. Introduction

First developed in the 1950s, fentanyl, a synthetic opioid that is 50 times stronger than heroin and 50-100 times more potent than morphine, has become a “miracle drug” used by pain specialists. Recently, it has become a major contributor to the opioid epidemic due to its widespread involvement in fatal and nonfatal overdose cases. About 39.5 million people lived with substance use disorders (SUD) in 2021^1^. Current reports indicate that about 52% of all overdose deaths involved illicit fentanyl^2^. Users who test positive for fentanyl and its analogs (F/FA) often do not realize their exposure to the substance^1^, which increases the likelihood of unintentional overdose events. The high potency of fentanyl and its rapid onset of action make fentanyl-laced street mixtures (e.g., heroin and fentanyl, methamphetamine and fentanyl) particularly lethal and hard to counteract in a timely fashion. Current medications used for the treatment of opioid use disorders (OUD) and overdose reversal agents are limited to agonists and antagonists of μ-opioid receptors (MORs).

One of the most promising strategies to curb OUD and the opioid epidemic is the development of vaccines against target opioids^3–5^. Active immunization induces the production of opioid-specific polyclonal antibodies that selectively bind the target drug, preventing it from crossing the blood-brain barrier to reach the brain where MORs and other opioid receptors are located. Thus, vaccines can attenuate opioid-induced behavior and pharmacological effects as well as toxicity associated with overdose^3^. Vaccines provide longer protection compared to small molecule therapeutics due to the long half-life of antibodies as well as the presence of antibody-producing plasma cells. Due to their selectivity, vaccines do not interfere with endogenous neuropeptides nor with medications for OUD^6, 7^.

As fentanyl itself is not immunogenic, a fentanyl analog must be conjugated to an immunogenic carrier protein or other foreign large molecule to create an antigenic motif or hapten that is recognized by hapten-specific B cells in the presence of cognate carrier-specific CD4^+^ T cells. We have previously reported the discovery and development of the conjugate F-CRM comprised of a fentanyl-based hapten containing a tetraglycine linker (F(Gly)_4_ or F) covalently conjugated to the genetically detoxified diphtheria toxin CRM_197_ ^6^. As subunit antigens are often too weak to induce a robust immune response by themselves, vaccine adjuvants are indispensable components to enhance vaccine immunogenicity and efficacy^3, 4^. In the context of anti-opioid vaccines, different adjuvant systems like aluminum salts^8^, toll-like receptor (TLR) 4 agonist monophosphoryl lipid A (MPLA) formulated in liposomes^9, 10^, TLR7/8 agonist INI-4001^11, 12^, TLR9 agonist CpG oligodeoxynucleotides (CpG ODNs) combined with Imject^®^ Alum containing aluminum hydroxide and magnesium hydroxide^13^, dsRNA^14^, heat-labile toxin of E. coli including dmLT and LTA1^15^, Freund’s adjuvant^16^, army liposome formulation (ALF)^17^, MF59^18^, Advax^TM^ ^19^ have been evaluated to produce optimal antibody titers. Other studies reported the use of immunomodulators targeting cytokines such as interleukin 4 (IL-4)^20, 21^. Results suggest that each vaccine formulation needs to be rationally designed, and the choice of adjuvants should be precisely tailored according to the nature of the drug-based hapten, the target immune response, and the patient population^22^. For instance, we previously reported that TLR4 agonists are not suitable for fentanyl vaccines (and possibly other opioid vaccines) due to competition of the TLR4 agonist with the F hapten at the TLR4^11^. TLR7/8 agonists may be particularly well suited for anti-opioid vaccines due to their ability to bias a T helper 1 cell (Th1) immune response and enhance antigen-specific IgG2a titers^21, 23^. Our team has designed a synthetic TLR7/8 agonist INI-4001 and evaluated its efficacy when combined with the lead anti-fentanyl vaccine candidate F-CRM. The combination of INI-4001 and Alhydrogel^®^ (alum) achieved a dose-sparing effect, significantly enhanced fentanyl-specific antibody titers and further reduced brain fentanyl concentration in rodents and porcine models compared to F-CRM adsorbed on alum^11, 12^. This formulation was shown to increase the mean polyclonal antibodies avidity for the fentanyl hapten compared to the vaccine formulated in alum alone^11^. Although aluminum salts have likely been the first adjuvant choice when designing a new vaccine for many years, vaccines formulated with aluminum salts cannot be filter sterilized, lyophilized, or frozen, and the administration route of approved aluminum-based vaccines, except for the Anthrax vaccine, is limited to intramuscular (IM) injection.

Inspired by INI-4001 efficacy and our goal to replace aluminum-based vaccine formulations, this study investigated the potential of liposomes containing INI-4001 as a viable alternative adjuvant to INI-4001/alum formulation. INI-4001 has two lipid tails, allowing the insertion of INI-4001 into the lipid bilayer of liposomes while displaying INI-4001’s hydrophilic active segment on the liposome’s surface^24^. The impact of liposome surface charges on F-CRM antigen adsorption and their correlation with *in vivo* efficacy was evaluated. To shed some light on the mechanism of addiction vaccines on innate immunity, we investigated the effect of the adjuvanted fentanyl vaccine on antigen-presenting cell subsets *in vitro*. We showed that cationic INI-4001 liposomes were as effective as INI-4001/alum adjuvanted vaccine in protecting rats against a fentanyl challenge and inducing the highest levels of activation and maturation on dendritic cells (DCs) relative to other INI-4001 liposomal formulations, while cationic INI-4001 liposomes were superior in inducing macrophage activation. Novel analytical ultracentrifugation methods also confirmed that F-CRM is most effectively incorporated into cationic liposomes, which explains this observation. These data support the use of INI-4001 cationic liposomal formulation as an adjuvant system for effective vaccines against fentanyl and other opioids, aiding in the treatment of OUD and the prevention of opioid overdose.

## 2. Materials and methods

### 2.1 Materials

Research-grade phospholipids and lipids such as 1,2-dioleoyl-sn-glycero-3-phosphocholine (DOPC), 1,2-dioleoyl-sn-glycero-3-phospho-(1’-rac-glycerol) (DOPG), 3ß-[N-(N’,N’-dimethylaminoethane)-carbamoyl]cholesterol hydrochloride (DC-Chol) and cholesterol (Chol) were purchased from Avanti® Polar Lipids (Irvine, CA, USA). GMP-grade *E. coli*-expressed diphtheria cross-reactive carrier protein-197 (EcoCRM) was purchased from Fina Biosolutions, LLC (Rockville, MD, USA). N-ethyl-N′-(3 dimethylaminopropyl) carbodiimide hydrochloride (EDAC), N-hydroxysuccinimide (NHS) and Fluorescein-5-Isothiocyanate (FITC ‘Isomer I’) were obtained from Sigma-Aldrich (St. Louis, MO, USA). Triethylamine (TEA), dimethyl sulfoxide (DMSO), sterile water for irrigation (WFI), assay buffers, and analytical grade reagents for HPLC were purchased through Thermo Fisher Scientific (Waltham, MA, USA). Alhydrogel^®^ (alum) was purchased from InvivoGen (San Diego, CA, USA). Amicon spin filters (MWCO 30 kDa) were obtained from Millipore, Merck (Burlington, MA, USA). Supor® PVDF syringe filters (0.8/0.22 µm) were purchased from PALL Inc. (Ann Arbor, MI, USA). The TLR7/8 agonist, INI-4001, was synthesized and processed to over 99% purity as previously described^11^. The fentanyl hapten containing a tetraglycine linker (F) and F-CRM conjugate were prepared following our previously published methods with some minor modifications^6, 25^.

### 2.2 Liposomal and aqueous formulations of INI-4001

To investigate the effect of differently charged liposomes loaded with INI-4001 on the efficacy of the fentanyl vaccine, INI-4001 anionic liposome DOPG/Chol, INI-4001 neutral liposome DOPC/Chol, and INI-4001 cationic liposome DOPC/DC-Chol were prepared. Briefly, liposomes were prepared using a thin-film hydration method with modification^24^. Stock solutions of 125 mM phospholipids DOPG and DOPC were made in chloroform. Cholesterol and DC-Chol stock solutions at 125 mM were prepared in chloroform. Phospholipid and cholesterol at a 2:1 molar ratio were combined in a 4 mL Covaris vial. INI-4001 was dissolved in chloroform and added to the phospholipid:cholesterol mixture at a weight ratio of INI-4001:phospholipid of 1:20 (w/w) and targeted 2.0 mg/mL in the final liposome formulations (**Table 1**). After mixing at room temperature (RT) for 5 minutes, the organic solvent was removed using a rotary evaporator at 250 mBar for 3 minutes, followed by 50 mBar for 10 minutes at 40 °C to form a thin film in the vial (Hei-VAP Expert HL, Heidolph Instruments GmbH & Co. KG, Schwabach, Germany). Dried thin films were kept under vacuum at RT overnight to remove traces of solvent. Afterward, thin films were hydrated in 50 mM Tris, 75 mM NaCl buffer pH 7.4 assisted by a focused-ultrasonicator with a Covaris S2 (Woburn, MA, USA) for 45 to 60 minutes to reduce the droplet size to below 80 nm. Liposomes were sterile filtered by syringe filters using PVDF membrane 0.8/0.22 µm (Supor^®^ membrane, PALL Inc., MI, USA).

**Table 1.** Composition of charged liposome formulations.

| Liposomes | Phospholipid (50 mM) | Chol/DC-Chol (25 mM) | INI-4001 (µg/mL) | Surface charge |
| --- | --- | --- | --- | --- |
| DOPG/Chol | DOPG | Chol | - | Anionic |
| DOPC/Chol | DOPC | Chol | - | Neutral |
| DOPC/DC-Chol | DOPC | DC-Chol | - | Cationic |
| INI-4001/DOPG/Chol | DOPG | Chol | 1868.085 | Anionic |
| INI-4001/DOPC/Chol | DOPC | Chol | 1774.164 | Neutral |
| INI-4001/DOPC/DC-Chol | DOPC | DC-Chol | 1822.668 | Cationic |

The aqueous 4001 formulation was used as a positive control to compare the activity of newly made INI-4001 encapsulated liposomes *in vivo*. Briefly, INI-4001 was weighed in a glass vial, and an adequate amount of aqueous buffered vehicle comprising 50 mM TRIS 1.5% glycerin and 0.1% Tween 80 pH 7.5 was added. The mixture was homogenized using a high shear homogenizer (Silverson L5MA, Silverson Machines Ltd, MA, USA) at 10,000 rpm for 20 minutes until the particle size was below 200 nm. The aqueous formulations were sterile filtered using a 13 mm Millex GV PVDF filter with a pore size of 0.22 μm (MilliporeSigma, MA, USA). All the INI-4001 encapsulated liposomes and aqueous INI-4001 formulations were stored at 2-8 °C until further use.

### 2.3 Characterization of INI-4001 liposomal and aqueous formulations

The physicochemical properties of liposomes were characterized, including particle size, polydispersity index (PDI), zeta potential, pH, osmolality, and stability of particle properties at different storage conditions. The particle size, PDI, and zeta potential were measured by the dynamic light scattering (DLS) method using Zetasizer Nano-ZS (Malvern Analytical, Malvern, UK). For particle size and PDI determination, samples were diluted 50 times in WFI. Particle size was measured at 25 °C with backscatter (173°) detection mode and reported as Z.Avg diameter based on the intensity function. The particle size measurement also provided the uniformity and dispersity of the liposomes indicated by PDI. For zeta potential measurement, samples were diluted 50 times in 1 mM NaCl solution and measured using a folded capillary cell at 25 °C. The pH value of each liposome sample was determined using an Accumet AB150 pH meter (Thermo Fisher Scientific) and an InLab Microprobe (Mettler-Toledo, Columbus, OH, USA) calibrated with a three-point curve at pH values 4, 7 and 10. Osmolality was measured using a Wescor Vapro 5520 vapor pressure osmometer. All measurements were performed in triplicate.

INI-4001 was quantified by reverse phase-high pressure liquid chromatography (RP-HPLC) using a Waters Acquity Arc UHPLC separation system equipped with a Waters 2998 PDA Detector. Briefly, samples were diluted in THF:methanol mixture (9:1, v/v) and eluted on a CORTECS^®^ C18 2.7 µm 3.0 x 50 mm column (P/N: 186007400). Mobile phase A consisted of 0.5% of 1 M ammonium formate and 9% of methanol in 90.5% of water. Mobile B consisted of 0.5% of 1 M ammonium formate and 99.5% of methanol. Elution was obtained by using the following gradient steps of solvents A and B: 85:15 (A:B) for the initial 2 minutes, 100% B for the next 10 minutes, and 85:15 (A:B) for the final 3 minutes at a flow rate of 1 mL/minute. The absorbance of INI-4001 was measured at 280 nm, and its content was determined by interpolation from a six-point standard dilution series ranging from 50 to 800 µg/mL.

### 2.4 Vaccine formulation preparation for mouse and rat *in vivo* studies

Prior to the injection in mice or rats, F-CRM conjugates were buffer exchanged to 50 mM Tris 1.5% Glycerol buffer pH 7.4. For vaccines using INI-4001 liposomes, F-CRM conjugates were mixed with INI-4001 encapsulated liposomes. For vaccines using aqueous INI-4001 and alum combination, F-CRM conjugates were adsorbed to alum before mixing with aqueous INI-4001. The formulation was placed on a rotator for one hour to allow the complete adsorption of INI-4001 to the F-CRM/alum. The final dose was 5 µg F-CRM, 10 µg INI-4001, and 24 µg alum.

### 2.5 Determination of F-CRM and INI-4001 absorption on aluminum salts

F-CRM conjugates were incubated with alum on a rotator at RT for one hour to facilitate the adsorption of the anionic F-CRM protein conjugates onto the cationic alum particles. Following incubation, the F-CRM/alum mixture was centrifuged at 10,000 rpm for 10 minutes to induce the sedimentation of alum particles. The supernatant was collected and analyzed in triplicate for protein content using the bicinchoninic acid (BCA) protein assay, as described in the manufacturer’s instructions. Absorbance was measured at 562 nm using a microplate reader, and protein concentrations were quantified based on a calibration curve prepared with bovine serum albumin standards (62.5–2000 µg/mL). Similarly, INI-4001 was mixed with the F-CRM/alum complex and rotated at RT for one hour. The formulation mixture was then centrifuged at 10,000 rpm for 10 minutes, and the INI-4001 content in the supernatant was quantified using RP-HPLC. Details of the HPLC conditions are provided in Section 2.3.

### 2.6 Determination of F-CRM adsorption on INI-4001 liposome formulations using analytical ultracentrifugation

This technique was exploited to examine the interaction between F-CRM conjugates and differently charged liposomes by measuring the sedimentation velocities of proteins or particles.

For this study, a FITC labeled F-CRM control was prepared at 1.6 µM according to the manufacturer’s protocol with minor modifications. FITC was dissolved in DMSO to the final concentration of 10 mg/mL and added to the F-CRM protein solution at 1.5 mg/mL in 10 mM phosphate buffer pH 7.4 containing 250 mM sucrose. The final molar ratio of FITC:F-CRM in the reaction mixture was 30:1. The coupling reaction was allowed to proceed overnight at 4 °C, under light-protected conditions. Subsequently, FITC labeled F-CRM protein was purified by washing with fresh buffer using a 30 kDa MWCO Amicon spin filter until no detectable fluorescence was observed in the filtrate, as assessed by UV absorbance at 495 nm. The purified FITC F-CRM product was characterized for its FITC-to-protein molar ratio (F/P), which was determined to be 0.093 using the equation below:

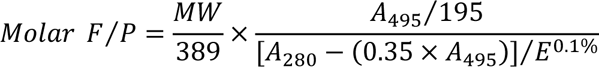

Where:

MW is the molecular weight of F-CRM protein,
389 is the molecular weight of FITC,
*A*_495_ and *A*_280_ are absorbance readings of FITC labeled F-CRM at 495 nm and 280 nm,
195 is the absorption *E*^0.1%^ of bound FITC at 490 nm at pH 13.0,
(0.35 × *A*_495_) is the correction factor due to the absorbance of FITC at 280 nm,
*E*^0.1%^ is the absorption at 280 nm of F-CRM protein at 1.0 mg/mL.

Cationic, anionic, and neutral INI-4001 liposomes were prepared at high and low concentrations by diluting the stock solutions 80- and 250-fold, resulting in final concentrations of 24 µg/mL and 8 µg/mL, respectively. FITC F-CRM:liposome mixtures were prepared with a final concentration of 1.6 µM FITC F-CRM and 35.2 µg/mL liposomes. All samples were diluted with 50 mM Tris 75 mM NaCl pH 7.4 buffer. The sedimentation velocity experiments of all controls and mixtures were measured using Beckman-Coulter analytical ultracentrifugation instruments at the Canadian Center for Hydrodynamics at the University of Lethbridge. The liposome controls were measured in an Optima AUC^TM^ using 1.2 cm epon standard 2 channel centerpieces, scanning in intensity mode at 227 nm and 273 nm with UV optics at 27,000 rpm for 15 hours at 20 °C. The FITC labeled F-CRM control and FITC F-CRM:INI-4001 liposome mixtures were measured in a Proteomelab XLA equipped with the Aviv fluorescence detector and measured in 3 mm titanium 2 channel centerpieces at 27,000 rpm and 20 °C for 12 hours. All samples were measured at the same photomultiplier voltage and gainset, so fluorescent yield is directly proportional to the measured intensity for all samples.

Intensity data was collected using the UltraScan data acquisition module^26, 27^. Fluorescence intensity data was collected with the Aviv AOS software. All experiments were analyzed with UltraScan. Models were fitted using the 2-dimensional spectrum analysis (2DSA) in a multi-step refinement process to remove time- and radially-invariant noise, as well as to determine the boundary conditions^28, 29^. Refined data was further analyzed with the parametrically constrained spectrum analysis (PCSA)^30^ and the van Holde Weischet analysis^31^, which generates diffusion-corrected integral sedimentation coefficient distributions.

### 2.7 Animal subjects and animal studies

Animal studies were carried out in OLAW and AAALAC-accredited *vivarii* in accordance with the University of Montana, the University of Washington, and the University of Minnesota’s IACUC guidelines for the care and use of laboratory animals. Animals were group-housed under a standard 12-hour light/dark cycle and were given food and water *ad libitum*. All experiments were conducted during the light cycle. Six-to eight-week-old female BALB/c mice (n = 8 per group) were obtained from Jackson Laboratory (Bar Harbor, ME). Eight weeks of male Sprague Dawley rats (n = 7 per group) were obtained from Envigo and allowed to acclimate for approximately one week before vaccination.

#### Immunization and blood collection

Mice and rats were vaccinated IM with formulations containing 5 µg F-CRM, 10 µg of INI-4001, and 24 µg alum on days 0 and 14 for mice and days 0, 21, 42 for rats. Vaccines were diluted in sterile 50 mM Tris, 1.5% Glycerol buffer pH 7.4 to a final volume of 50 µL (mice) or 100 µL (rats) per injection. Animals injected with buffer were used as a negative control. One week after the last vaccination, mice were terminally bled, and serum was collected via terminal cardiac puncture. In contrast, rats were bled via tail vein, and sera were used for antigen-specific IgG titers analysis.

#### Fentanyl hapten-specific antibody titers

Anti-fentanyl hapten IgG-specific titers were measured via ELISA-based assay as described previously^6,15^. Briefly, ninety-six well Costar assay plates (Corning 3690) were coated with 5 ng/mL of fentanyl hapten F-BSA conjugate in carbonate coating buffer overnight at 4 °C. The plates were washed five times with 1x PBS + 0.05% Tween-20 (PBS-T) before being blocked with 1% porcine gelatin-PBS pH 7.4 for 1 hour at RT. The blocking solution was removed, and serum samples were added and serially diluted in PBS-T across the plates at 1:200, followed by a 2-hour incubation period at RT. Subsequently, plates were washed and incubated overnight at 4 °C with the following horseradish peroxidase-conjugated secondary antibodies: Goat anti-rat IgG (1: 50,000 dilution), goat anti-mouse IgG-HRP (Jackson ImmunoResearch, West Grove, PA), goat anti-mouse IgG1-HRP or IgG2a-HRP (Alpha Diagnostic International, Inc., San Antonio, TX, catalog #40126 and 40127, respectively) for IgG subclass titers. All mice’s secondary antibodies were diluted at 1:30,000 in PBS-T. The following day, plates were developed using SIGMAFAST OPD (o-phenylenediamine dihydrochloride) substrate (Sigma-Aldrich, St. Louis, MO) and read at 492 nm on a Tecan Infinite M1000 PRO Microplate Reader.

#### Assessment of association rates of fentanyl hapten-specific antibodies

All biolayer interferometry experiments were conducted at 30 °C and 1000 rpm using an Octet R8 and black polystyrene 96-well plates (Evergreen Labware Products #290-8195-Z1F). Total non-specific IgG concentrations were quantitated for each individual animal using Octet ProA Biosensors (Sartorius 18-5010). Subsequently, a kinetics experiment was conducted using Octet SA Biosensors (Sartorius 18-5019). Following a baseline measurement (1x PBST (Thermo Scientific 28352); 180 seconds), pre-hydrated SA probes were loaded with F-biotin (0.1ug/mL in PBST; 120 seconds), and baseline was measured again (1x PBST; 180 seconds). The association of polyclonal sera (200 nM, 100 nM, and 50 nM as determined by total IgG quantitation) was measured for 300 seconds, followed by 600 seconds of dissociation. Apparent association rate constants (K_a_) were calculated from raw data in Sartorius Analysis Studio v13.0.1.35. K_a_ values from curves with R^2^>0.95 were plotted in GraphPad Prism v10.3.1. The detailed results from this experiment can be found in Supplement S.4.

#### In vivo efficacy in rats

To assess the efficacy of vaccination on fentanyl-induced antinociception, respiratory and cardiovascular depression, and drug distribution, rats were challenged with a cumulative dose of 0.3 mg/kg fentanyl delivered SC through three sequential doses of 0.1 mg/kg. Prior to fentanyl administration, baseline arterial oxygen saturation (SaO_2_), heart rate, and breath rate were measured using a MouseOx Plus pulse oximeter (STARR Life Sciences Corp., Oakmont, PA) via neck collar for at least 1 minute to ensure stable readings. Immediately after oximetry measurements, baseline nociception was assessed by placing rats on a 54 °C hotplate and measuring the latency to perform hind paw licking, withdrawal or shaking, or jumping, with a 30 seconds maximum cutoff to prevent tissue damage. Rats then received 0.1 mg/kg fentanyl every 17 minutes to achieve cumulative dosing of 0.1, 0.2, and 0.3 mg/kg. Fifteen minutes after each fentanyl injection, antinociception and oximetry were measured for a total of 45 minutes. Following the last antinociception and oximetry measurements, rats were euthanized via CO_2_ inhalation, and blood and brain were collected for analysis of fentanyl concentration by liquid chromatography coupled with mass spectrometry (LC-MS) as previously described^6^.

### 2.8 Flow cytometry-based *in vitro* maturation and activation of macrophages and dendritic cells (DCs) assay

#### Cell lines

RAW 264.7 macrophage and JAWS II DC cell line (CRL-11904) were purchased from the American Type Culture Collection (ATCC). RAW 264.7 cells were maintained in Dulbecco’s modified Eagle’s medium (DMEM, Gibco, Carlsbad, CA) with 10% fetal bovine serum (FBS, Gibco, Carlsbad, CA) and 1% penicillin-streptomycin (Sigma-Aldrich). JAWS II DC cells were maintained in Alpha minimum essential medium with 5 ng/mL murine GM-CSF (ThermoFisher PMC2015), and 20% fetal bovine serum.

#### Cell activation

Macrophages and DC cells were plated at 0.7 × 10^6^ and 1 x 10^6^ cells/mL, respectively, in 6-well cell culture plates at 1 mL per well in complete medium (10% FBS for RAW264.7 and 20% for JAWS II) and allowed to adhere to the plate surface overnight. The next day, cells were serum starved for 6-12 hours in 0.1% FBS before adding stimulants. Fifty µL of vaccines were added to each well to achieve a final concentration of 5 µg/mL of F-CRM conjugate, 24 µg/mL of alum adjuvant, and 10 µg/mL of INI-4001, followed by 24 hours of incubation at 37 °C with 5% CO_2_. Cells incubated with medium only were used as a negative control. At the end of the incubation period, cells were harvested by gently scrapping (RAW264.7) or adding 0.5 mL of Trypsin-EDTA (0.25%), phenol red (Gibco, Carlsbad, CA, USA) followed by 0.5 mL of FBS to stop the digestion (JAWS II). All vaccine formulations were tested in triplicates.

#### Flow cytometry analysis

Cells were harvested and washed with 1X PBS followed by staining with Zombie NIR™ Fixable Viability Kit (1:500, BioLegend, San Diego, CA, USA) for 20 minutes at RT. Cell suspensions were washed using FACS buffer (1X PBS supplemented with 2% FBS), treated with mouse FcR Blocking Reagent (Miltenyi Biotech, Bergisch Gladbach, Germany), and then stained for 20 minutes at 4 °C with the following rat anti-mouse antibodies from BioLegend (San Diego, CA): APC anti-mouse CD86 Antibody (105011), Brilliant Violet 421™ anti-mouse I-A/I-E Antibody (107631), FITC anti-mouse CD40 Antibody (124607). Cells were then fixed and permeabilized using the BD Cytofix/Cytoperm Kit (BD Biosciences) for 20 minutes at 4 °C, followed by the addition of BD Perm/Wash Buffer (1X). Intracellular staining of RAW264.7 was performed by incubating with PE anti-Nos2 (iNOS) Antibody. Cells were acquired using a BD FACSymphony™ A1 Cell Analyzer, and data were analyzed using FlowJo software (BD Biosciences, San Jose, CA, USA).

### 2.9 Statistical analysis

Statistical analyses were performed using GraphPad Prism version 9.0 (GraphPad Software, San Diego, CA, USA). For parametric analysis, group values were confirmed for normality using the D’Agostino-Pearson omnibus normality test. Antibody titers and brain and serum fentanyl concentration were compared using one-way ANOVA with either Dunnett’s test or Tukey’s multiple comparisons post hoc test. Hot plate and oximetry data over time were compared using two-way ANOVA or mixed-effects analysis with Tukey’s multiple comparisons post hoc test. Median fluoresce intensity (MFI) data were analyzed via Brown-Forsythe and Welch’s ANOVA test.

## 3. Results

### 3.1 Characterization of INI-4001 liposomal formulations

In this study, three different charged liposomal formulations were prepared with phospholipid and lipid concentrations of 50 mM and 25 mM, respectively (**Table 1**).

**Table 2** and **Figure 1** summarize the particle size, PDI, and zeta potential of all liposome formulations. The liposomes ranged in size from 30 to 60 nm and exhibited a PDI below 0.3, which is desirable for sterile filtration and long-term colloidal stability. The incorporation of INI-4001 into anionic DOPG/Chol and cationic DOPC/DC-Chol liposomes had minimal effect on particle size and size distribution; however, in neutral DOPC/Chol liposomes, a reduction in particle size was observed. The inclusion of INI-4001 increased the negative surface charge of anionic liposomes and reduced the positive surface charge of cationic liposomes, while maintaining the zeta potential of neutral liposomes within the neutral range. In contrast, the addition of F-CRM conjugates to INI-4001 liposomes significantly altered both the size and zeta potential across all liposome formulations.

**Figure 1.**
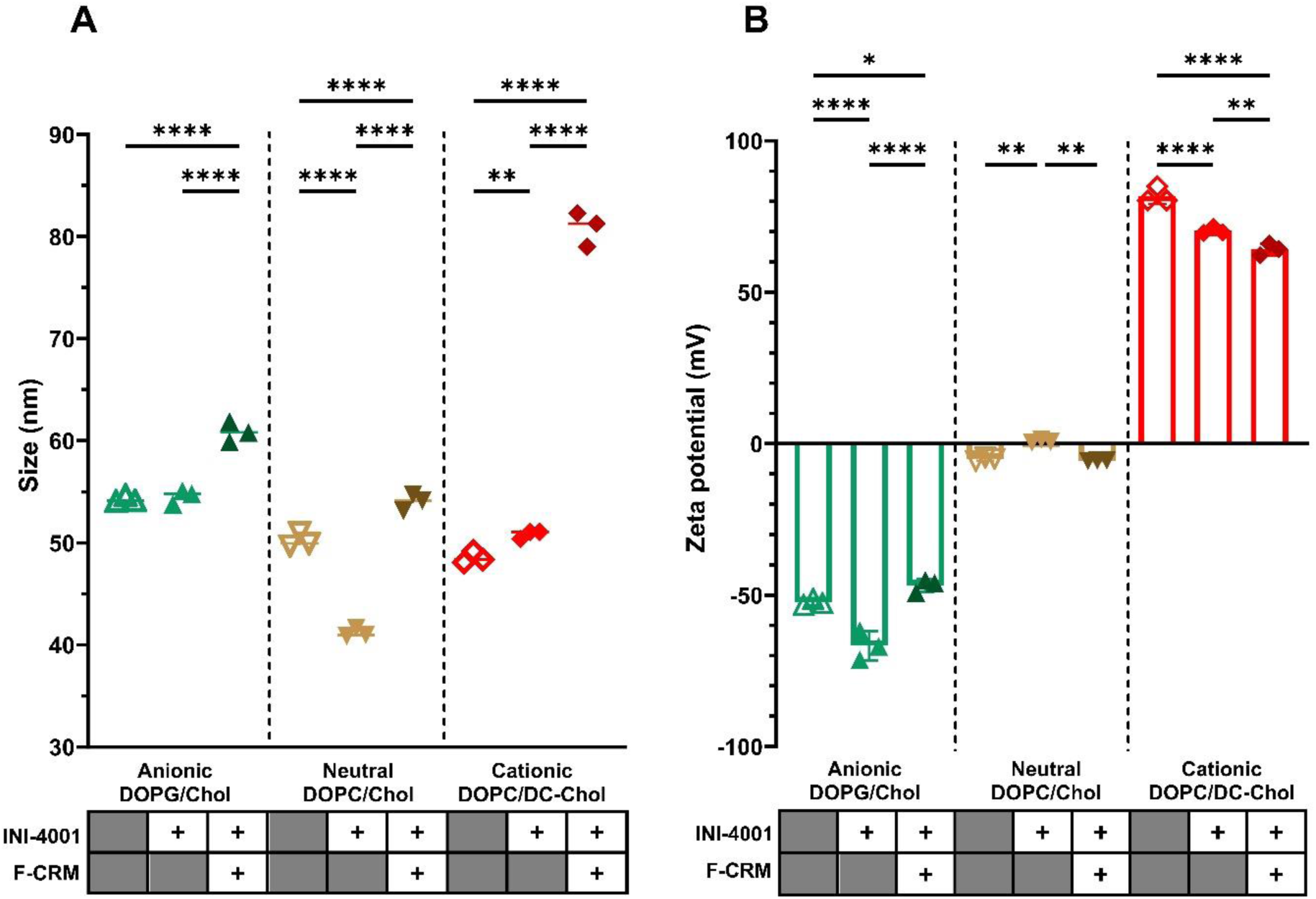
Dynamic light scattering characterization. (A) Size and (B) zeta potential of differently charged liposomes without INI-4001, with INI-4001 and F-CRM. Statistical analysis was conducted using two-way ANOVA with Tukey’s multiple comparisons. *p ≤ 0.05, **p ≤ 0.01, ***p ≤ 0.001, ****p ≤ 0.0001. Data expressed as mean ± standard deviation (n = 3).

**Table 2.**
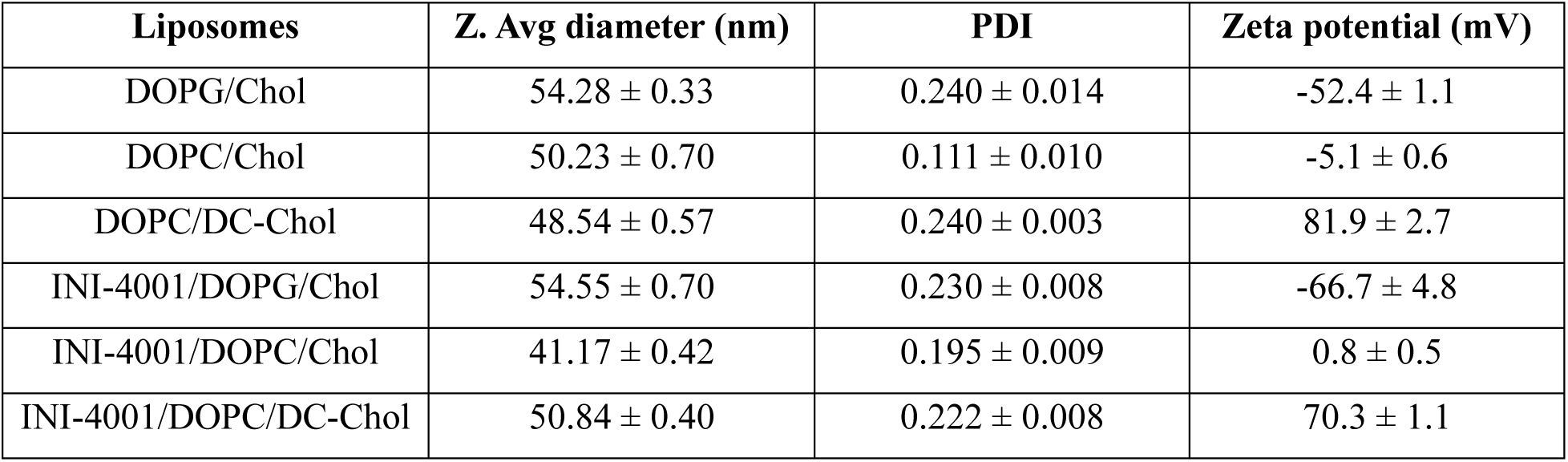
Size, PDI, and zeta potential of liposomal formulations with and without INI-4001. Data expressed as mean ± standard deviation (n = 3)

All liposomal formulations were isotonic and had pH values within the acceptable range for parenteral administration. HPLC data showed a high recovery of INI-4001 (Supplementary S.1).

### 3.2 F-CRM and INI-4001 adsorbed to alum

The F-CRM conjugate, an anionic monomeric protein with a molecular weight of 64.1 kDa, was characterized using gel electrophoresis and Matrix-Assisted Laser Desorption/Ionization Time-of-Flight (MALDI-TOF) mass spectrometry as previously described^32^. Due to its net negative charge, the F-CRM conjugate was expected to interact strongly with positively charged alum through ionic interactions. This hypothesis was validated by assessing the adsorption of F-CRM onto alum. The evaluation involved centrifugation of the F-CRM and alum mixture to pellet the alum particles, followed by quantification of unbound protein in the supernatant using the BCA assay. The results indicated nearly complete adsorption (∼100%) of F-CRM onto alum (data not shown).

Similarly, the adsorption of INI-4001 onto the F-CRM/alum complex was analyzed by quantifying the supernatant after centrifugation of the INI-4001 + F-CRM/alum mixture at 10,000 rpm for 10 minutes. HPLC analysis revealed that nearly 100% of INI-4001 was adsorbed after mixing the INI-4001 + F-CRM/alum for 45 minutes (Figure 2).

**Figure 2.**
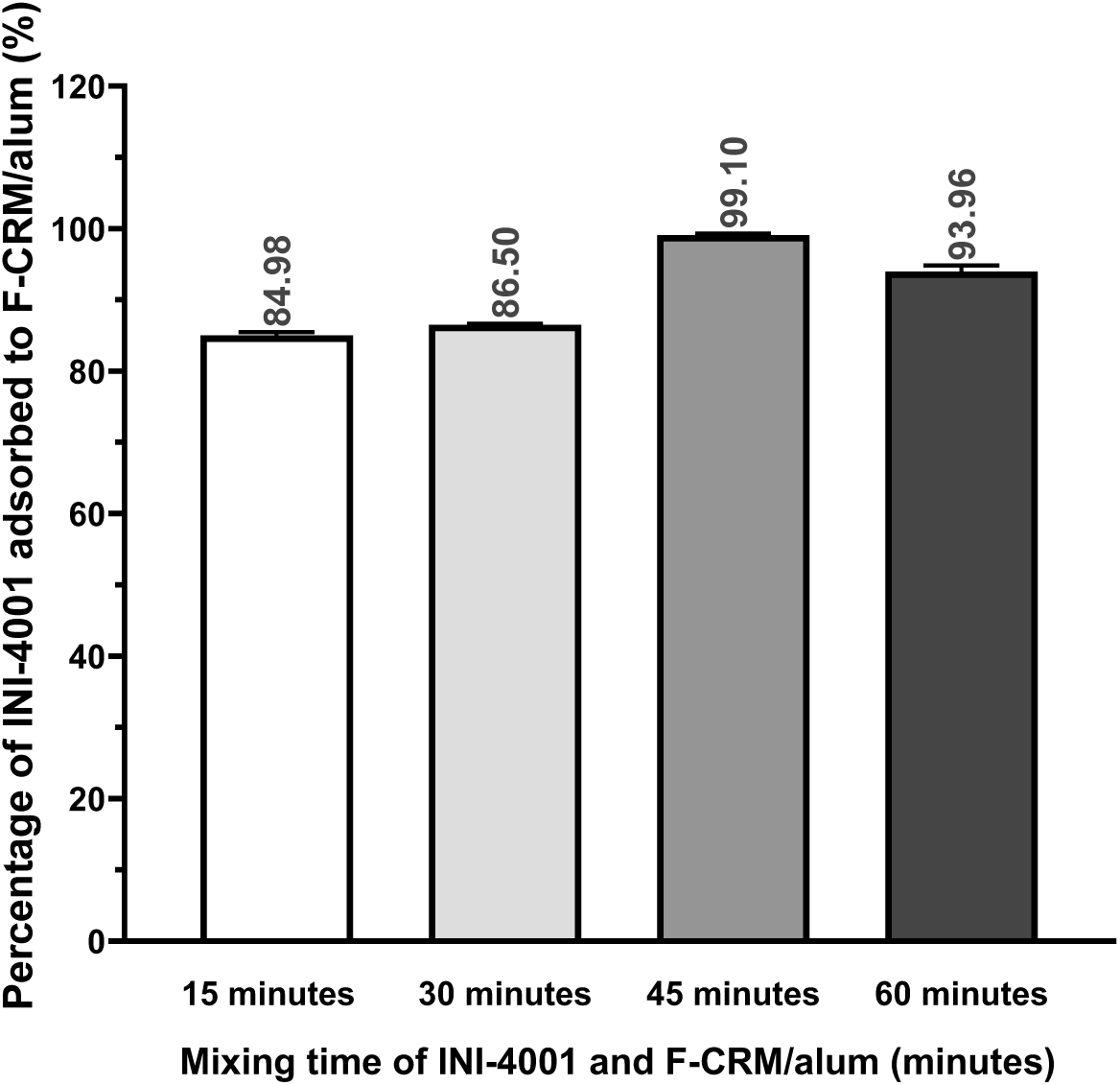
Time-dependence adsorption of INI-4001 to F-CRM/alum. Aliquots from INI-4001 + F-CRM/alum mixture were taken after mixing 15, 30, 45 and 60 minutes. Samples were centrifuged at 10,000 rpm for 10 minutes. The supernatants were quantified for INI-4001 content by HPLC. Data expressed as mean ± standard deviation (n = 3).

### 3.3 Cationic liposomes can strongly bind conjugate antigens

FITC was conjugated to the F-CRM protein at a F/P molar ratio of 0.037. This low F/P ratio was selected to minimize any potential alteration of the physicochemical properties of the F-CRM protein, which could influence its adsorption behavior on liposomal surfaces. The impact of FITC labeling on F-CRM was evaluated by comparing the size, zeta potential, and molecular weight of FITC F-CRM to the F-CRM protein. Insignificant differences were observed, as confirmed by size and zeta potential measurements by DLS, and molecular weight analysis via MALDI-TOF, indicating that the labeling process did not substantially affect the properties of the F-CRM protein (Supplementary S.2).

The interaction between the FITC labeled F-CRM conjugate and differently charged INI-4001 liposomes was further evaluated by the analytical ultracentrifugation. The sedimentation velocity of the FITC F-CRM and liposome mixtures was compared to that of the FITC F-CRM conjugate alone as a control. The homogeneity of the sample is assessed by comparing the sedimentation distributions generated by 2DSA-IT models. A high value of the sedimentation coefficient indicates large or high-density particles sedimenting early during centrifugation; a small value indicates lower-density or smaller particles sedimenting later during centrifugation, while a negative value indicates floating particles that have a density value smaller than that of the dispersing buffer. The binding of protein to liposomes would increase the overall density of the complex, leading to a right shift in the sedimentation profiles on van Holde-Weischet plots.

Characterization of the FITC labeled F-CRM control showed a homogeneous monomer. A pseudo-3D plot of the PCSA model is shown in Figure 3, and the molar mass integration results are shown in **Table 3**. The model reports an average molecular weight of 63.3 kDa, which is in excellent agreement with the sequence calculated molecular weight of 64.1 kDa for a F-CRM monomer.

**Figure 3.**
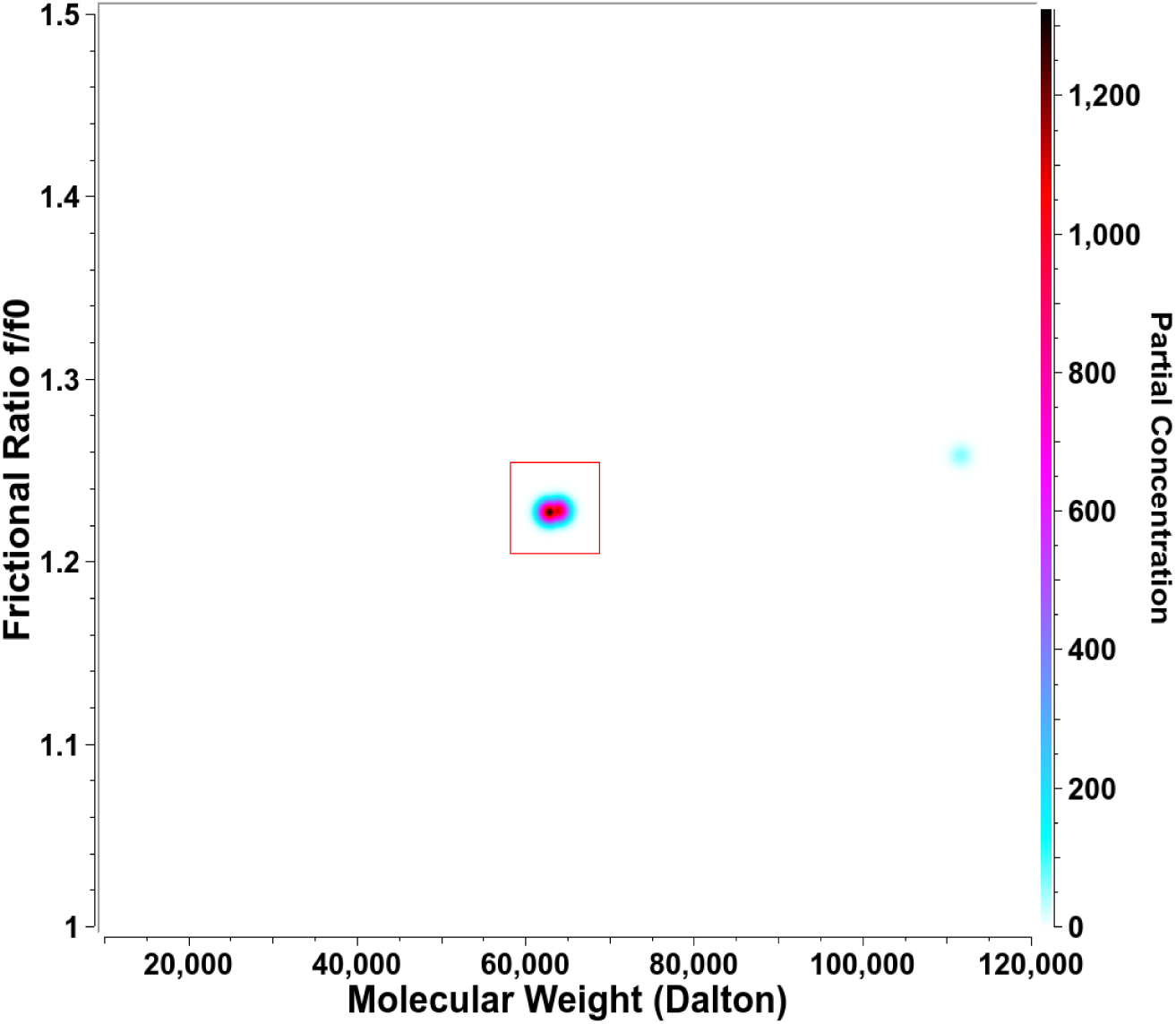
Pseudo-3D plot of FITC labeled F-CRM control using a PCSA model. A single species is observed, see signal integration in Table 3.

**Table 3.** FITC labeled F-CRM PCSA model integration.

|  |  |
| --- | --- |
| Measured MW | 63.269 kDa |
| Sequence MW | 64.127 kDa |
| Sedimentation coefficient | $4.64 \times 10^{-13}$ S |
| $f/f_0$ | 1.23 |

Raw fluorescence data is shown in Figure 4, highlighting the various degrees of quenching observed when the FITC labeled F-CRM interacts with differently charged liposomes. The cationic INI-4001/DOPC/DC-Chol liposomes appeared to be significantly quenched compared to the control, yielding the least signal (∼200 intensity counts) compared to the FITC F-CRM control (∼3,400 intensity counts). Except for the quenched cationic liposome sample, a similar baseline offset of ∼250 counts is observed in all samples, which indicates a small amount of residual free FITC dye in the FITC labeled F-CRM control.

**Figure 4.**
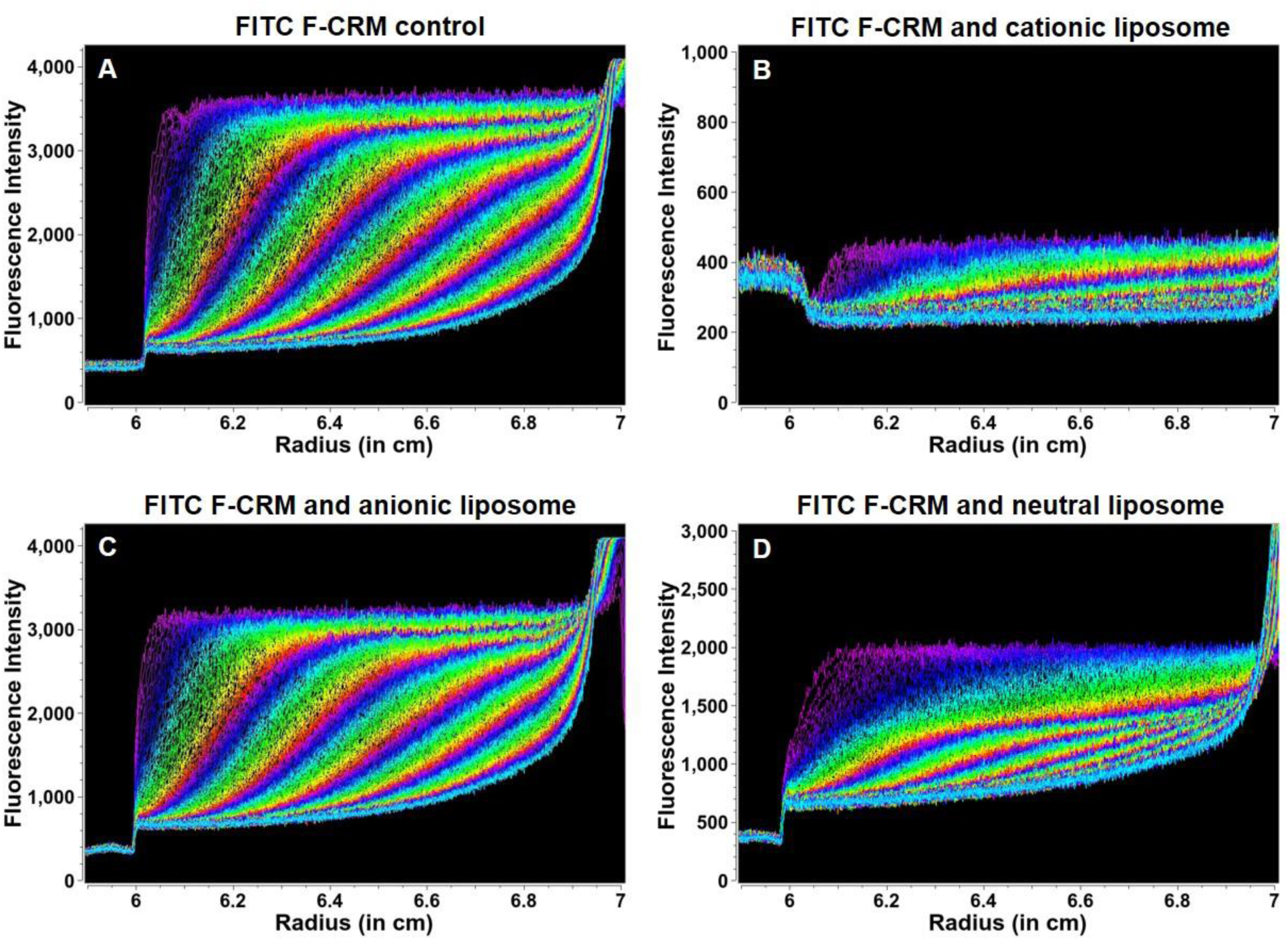
Raw fluorescence data. Analysis of FITC labeled F-CRM control (A) and its mixtures with cationic INI-4001/DOPC/DC-Chol liposome (B), anionic INI-4001/DOPG/Chol liposome (C), and neutral INI-4001/DOPC/Chol liposome (D). Raised baseline indicated free unsedimenting FITC dyes. Differences in overall intensity values with various liposomes suggest various levels of quenching in the presence of differently charged liposomes.

Integral sedimentation distributions for controls and mixtures are shown in Figure 5. Comparison of the high and low concentrations of the three liposomes resulted in virtually identical sedimentation coefficient distributions for each liposome type (Figure 5A). Cationic and neutral liposome controls, measured with UV detection, have a similar distribution from 1-50 S, while the anionic liposomes are distributed from 1-110 S (Figure 5A). Sedimentation profiles of the FITC F-CRM:liposome mixtures suggest that FITC F-CRM only binds to cationic and neutral liposomes (Figure 5B). Anionic INI-4001/DOPG/Chol liposomes mixed with FITC F-CRM show no difference between the mixture and the FITC F-CRM control (Figure 5C). Cationic INI-4001/DOPC/DC-Chol liposomes mixed with FITC F-CRM show a sedimentation distribution ranging between 5-30 S, and 80-90% of the FITC F-CRM signal sediments faster than the FITC F-CRM control, indicating liposome integration (Figure 5D). Neutral INI-4001/DOPC/Chol liposomes mixed with FITC F-CRM show a sedimentation distribution ranging between 5-40 S, but only 60-70% of the FITC F-CRM signal sediments faster than the FITC F-CRM control (Figure 5E).

**Figure 5.**
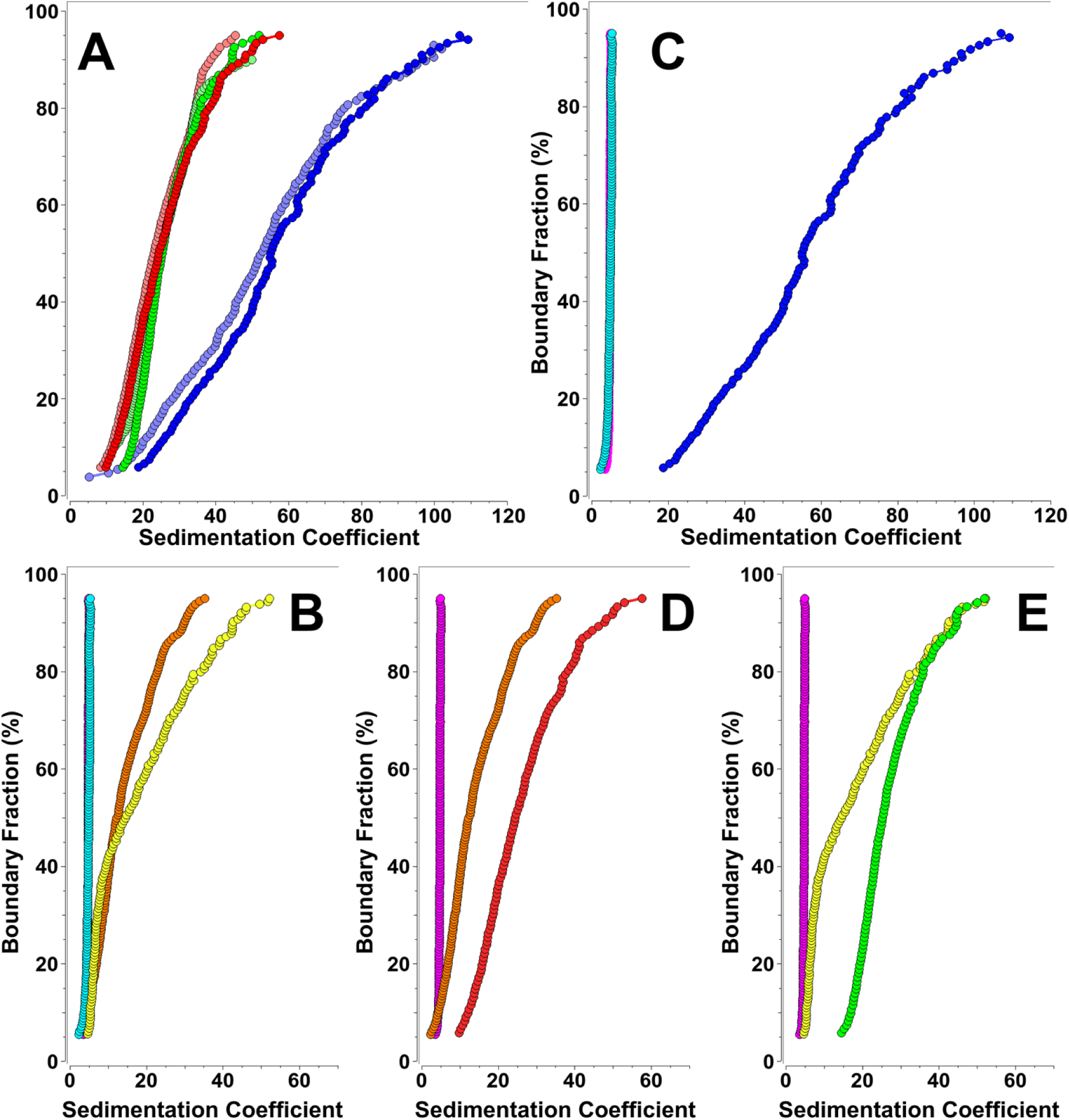
Analytical ultracentrifugation analysis of FITC F-CRM incorporation into anionic, neutral and cationic liposomes. (A) Integral s-value distributions of empty liposome controls measured with a UV detector. Red: cationic high concentration, light red: cationic low concentration. Blue: anionic high concentration, light blue: anionic low concentration. Green: neutral high concentration, light green: neutral low concentration. (B) Comparison of integral s-value distributions for all fluorescent samples and mixtures. FITC F-CRM control (magenta), and FITC F-CRM:liposome mixtures (cyan: anionic, yellow: neutral, orange: cationic). (C) Comparison of integral s-value distributions for FITC F-CRM control (magenta), anionic liposome control (blue), and FITC F-CRM:anionic liposome mixture (cyan). (D) Comparison of integral s-value distributions for FITC F-CRM control (magenta), cationic liposome control (red), and FITC F-CRM:cationic liposome mixture (orange). (E) Comparison of integral s-value distributions for FITC F-CRM control (magenta), neutral liposome control (green), and FITC F-CRM:neutral liposome mixture (yellow).

### 3.4 F-CRM adjuvanted with cationic liposomes induced robust immunogenicity and a Th1-biased response in mice

The effects of F-CRM and INI-4001 liposomal formulations on the production of fentanyl hapten-specific antibody titers were tested in an immunization study in BALB/c mice. All liposome formulations were compared to F-CRM/alum/INI-4001 as a positive control. Mice were vaccinated twice, two weeks apart, and serum was collected one week after the second immunization via terminal cardiac puncture and used to measure fentanyl hapten-specific antibody titers of IgG, IgG1, and IgG2a isotypes.

As shown in Figure 6, serum from mice vaccinated with F-CRM plus the cationic INI-4001/DOPC/DC-Chol liposomes showed comparable total IgG titer as INI-4001/alum adjuvanted control. Positively charged INI-4001/DOPC/DC-Chol liposomes surpassed neutral INI-4001/DOPC/Chol liposomes and anionic INI-4001/DOPG/Chol liposomes in triggering significantly higher fentanyl hapten-specific antibody responses. When liposomes were admixed with alum, we detected no significant differences in total IgG titer compared to liposomes without alum (Supplement S.3), suggesting that alum is not essential for liposome immunogenicity. As previously reported, adding INI-4001 to F-CRM/alum significantly enhanced the antibody’s avidity to F hapten^11^, we assessed whether cationic INI-4001/DOPC/DC-Chol liposomes modulate fentanyl hapten-specific polyclonal antibodies avidity compared to INI-4001/alum control using OctetRed biolayer interferometry (BLI). The results suggest that cationic liposomes enhance the immunogenicity of F-CRM while maintaining comparable antibody avidity to INI-4001/alum control (Supplement S.4). We then investigated IgG subtypes IgG1 and IgG2a as a representation of Th2- and Th1-type responses, respectively. The INI-4001/alum formulation demonstrated significantly higher anti-fentanyl hapten serum IgG1 titers than the liposome adjuvanted vaccines. The cationic INI-4001/DOPC/DC-Chol liposomes outperformed the neutral and anionic liposomes and INI-4001/alum for their ability to induce IgG2a titers. The ratio of IgG1 to IgG2a of fentanyl hapten-specific antibodies was significantly lower in liposome groups compared to the INI-4001/alum adjuvanted group, demonstrating a biased Th1 response in the former compared to a more balanced Th1/Th2 response in the latter (Figure 7).

**Figure 6.**
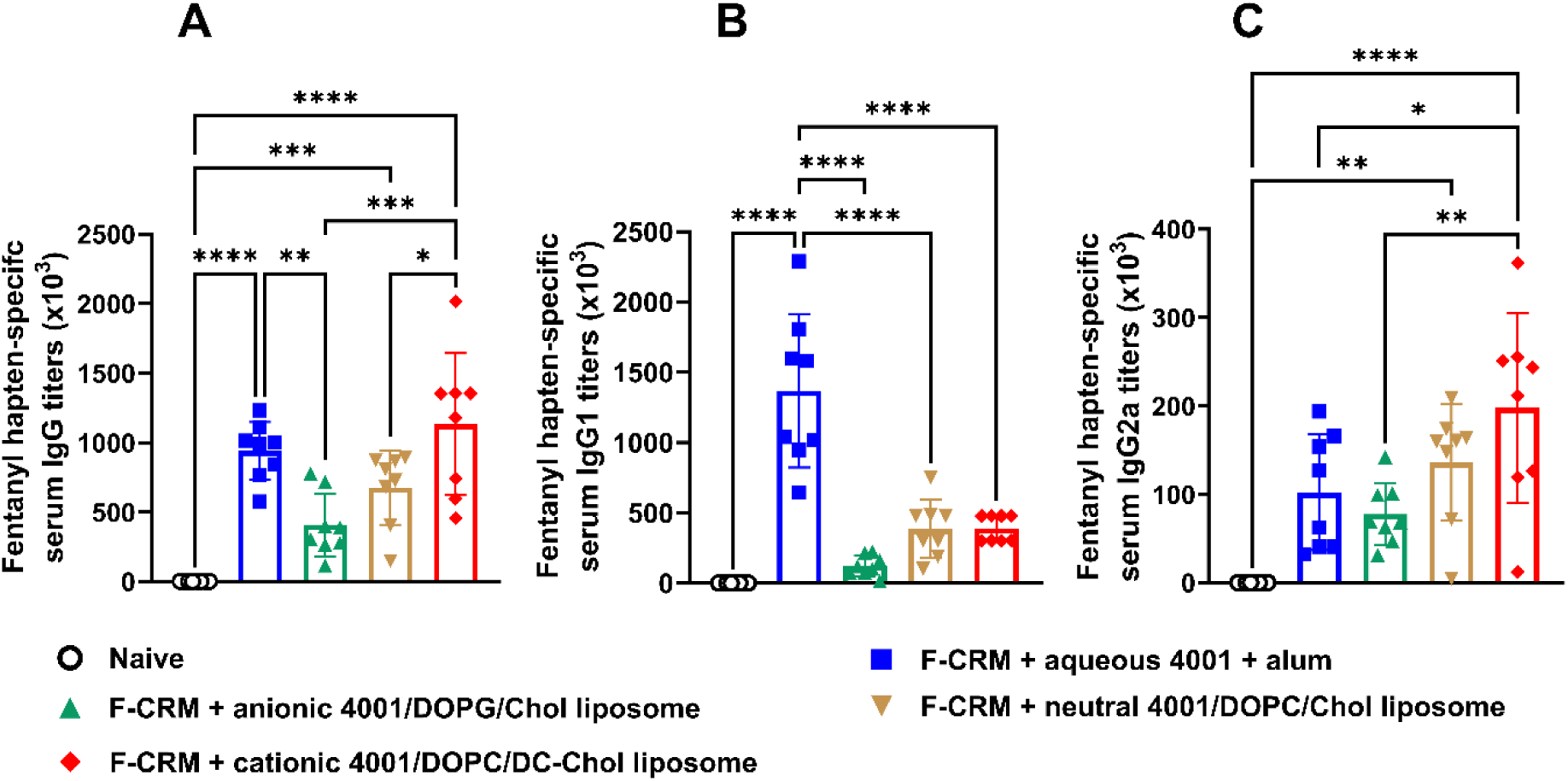
The immunogenicity and immunoglobulins (Ig) subclass when adjuvanting F-CRM with INI-4001 liposomes in mice. Six- to eight-week-old female BALB/c mice (n = 6-8) were vaccinated on days 0 and 14, IM, with 5 µg F-CRM plus 10 µg INI-4001 and 22.5 µg alum. The serum was collected on day 21. Anti-fentanyl hapten IgG (A), IgG1 (B), and IgG2a (C) antibody titers were measured via ELISA. Statistical analysis was conducted using one-way ANOVA and Tukey’s multiple comparisons. *p ≤ 0.05, **p ≤ 0.01, ***p ≤ 0.001, ****p ≤ 0.0001; color of asterisks indicates comparison group.

**Figure 7.**
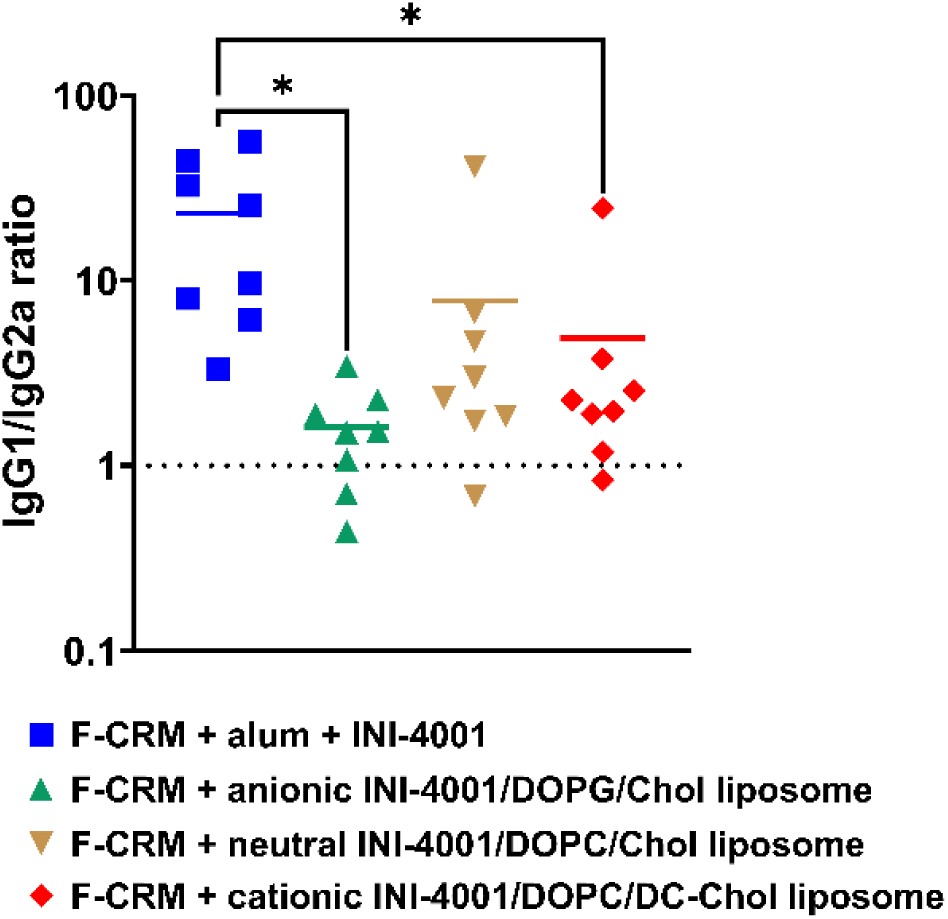
Antibody titer subclass ratio. Six- to eight-week-old female BALB/c mice were vaccinated on days 0 and 14, IM, with 5 µg F-CRM plus 10 µg INI-4001 and 22.5 µg alum. The serum was collected on day 21. Anti-fentanyl hapten IgG, IgG1, and IgG2a antibody concentrations were measured via ELISA. IgG1 and IgG2a ratios were calculated to examine Th2- and Th1-polarization changes. Statistical analysis was conducted using one-way ANOVA and Tukey’s multiple comparisons. *p ≤ 0.05; color of asterisks indicates comparison group. Data are shown as mean ± SD.

### 3.5 F-CRM adjuvanted with cationic liposomes induced robust antibody response and protected against fentanyl challenge in rats

We next evaluated the efficacy of F-CRM and INI-4001 liposomes in blocking fentanyl-induced pharmacological and side effects in Sprague Dawley rats. Rats received three vaccinations at 3 weeks intervals, and one week after the third vaccination, fentanyl hapten-specific antibody titers were analyzed using ELISA. Immunization with the cationic INI-4001/DOPC/DC-Chol liposomes induced the highest IgG antibody titers with a statistically significant difference (p = 0.0334) when compared to the lead formulation F-CRM/alum/INI-4001 (the second most effective) (Figure 8A). Consistent with murine data, the anionic INI-4001/DOPG/Chol liposome also showed poor immunogenicity in rats. Two weeks after the last vaccination, animals were challenged with 0.1 mg/kg fentanyl SC every 15 minutes to a total of 0.3 mg/kg cumulative fentanyl dose, and the antinociception (Figure 8D), oxygen saturation (Figure 8E), and heart rate (Figure 8F) were measured before drug administration and 15 minutes post each fentanyl injection. Fifteen minutes after the last fentanyl dose, brains and serum were collected to measure fentanyl concentrations (Figure 8B and C). Analysis of fentanyl distribution on vaccinated animals revealed that F-CRM formulated with the cationic INI-4001/DOPC/DC-Chol liposomes was as effective as F-CRM/alum/INI-4001 in sequestering fentanyl in the serum, as evidenced by the high drug concentration in serum (Figure 8B) and the lowest brain concentration (Figure 8C). The fentanyl vaccine adjuvanted with anionic or neutral liposomes was the least effective in blocking fentanyl distribution to the brain. In evaluating vaccine-induced immune responses in protecting against fentanyl-triggered antinociception and respiratory and cardiovascular depression, both the F-CRM/alum/INI-4001 and F-CRM + cationic INI-4001/DOPC/DC-Chol liposomes completely blocked fentanyl-induced antinociception (Figure 8D) and prevented fentanyl induced drop in oxygen saturation (Figure 8E) and heart rate (Figure 8F). Together, both mice and rat studies demonstrate the potent adjuvanticity of cationic liposomes containing TLR7/8 agonist (INI-4001) in enhancing the immunogenicity and efficacy of the anti-fentanyl vaccine.

**Figure 8.**
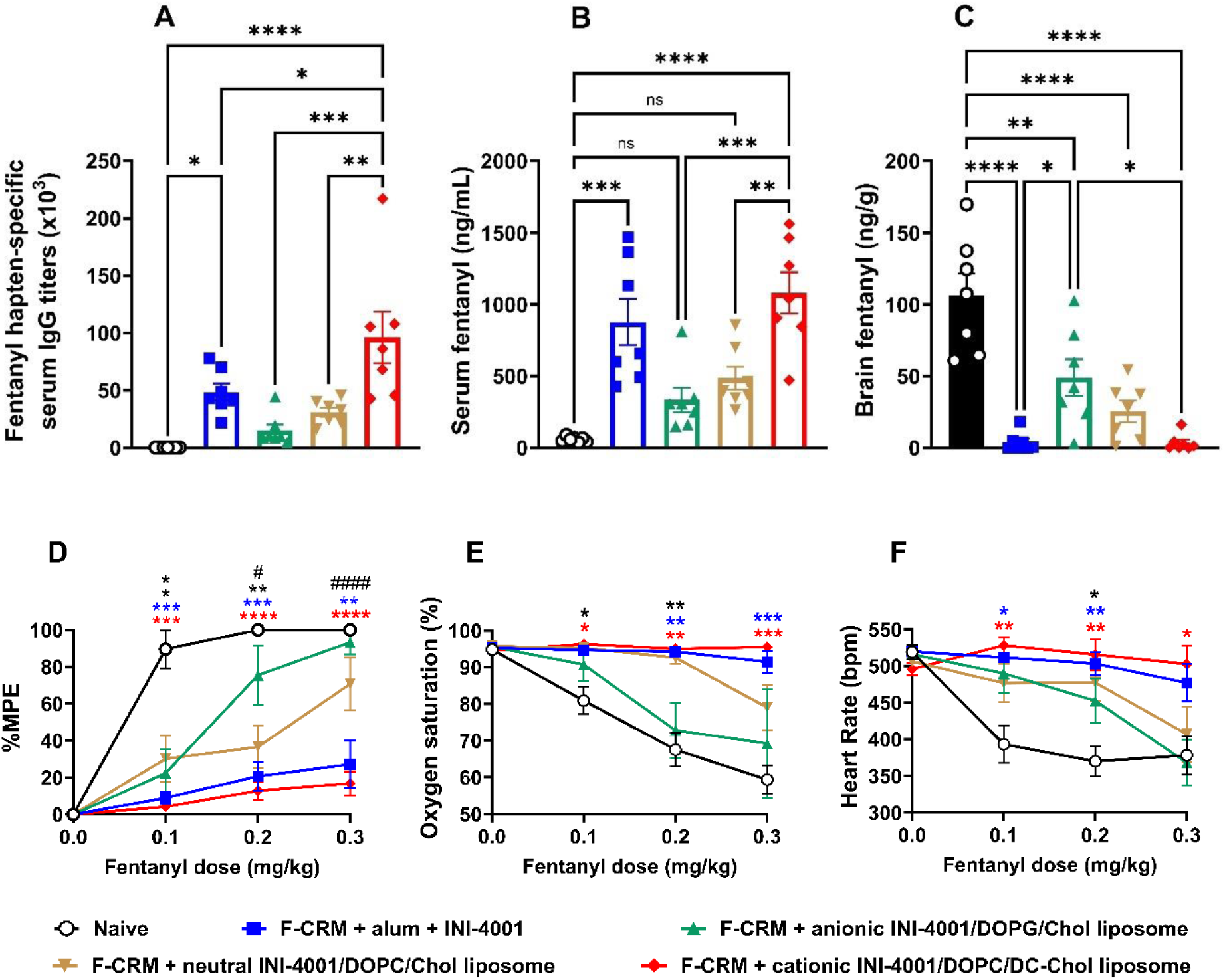
Immunogenicity and *in vivo* efficacy of vaccination against fentanyl with INI-4001 liposomal formulations in rats. Male Sprague Dawley rats (n = 7 per group) were vaccinated IM with F-CRM with INI-4001/alum or various INI-4001 liposomal formulations every 3 weeks for 3 vaccinations. Rats were bled via the tail vein one week after the third vaccination to analyze fentanyl hapten-specific antibody titers (A). Rats were subsequently challenged s.c. with 0.1 mg/kg every 15 minutes, and rats were tested 15 minutes post-injection for antinociception (D), respiratory depression (E), and bradycardia (F). Fifteen minutes after the last fentanyl dose, rats were euthanized, and brain and serum were collected to analyze fentanyl distribution via LC-MS. Fentanyl concentration was quantified in serum (B) and brain (C). Data were analyzed using one-way ANOVA with Tukey’s multiple comparisons post hoc test (A-C), two-way ANOVA with Tukey’s multiple comparisons post hoc test (D) or via mixed-effects analysis (E & F). Statistical symbols: **** or #### p < 0.0001, *** p < 0.001, ** p < 0.01, * or # p < 0.05. In D-F: * indicates significance compared to control, while # indicates significance between F-CRM + anionic INI-4001/DOPG/Chol liposomes vs. F-CRM + cationic INI-4001/DOPC/DC-Chol liposomes. Data are shown as mean ± SEM.

### 3.6 Cationic INI-4001 liposome induces the highest levels of the co-stimulatory markers CD86 and CD40 on DCs and macrophages

Our observation that INI-4001 liposomes with different surface charges induce quantitatively distinct *in vivo* responses suggests that these differences may be mediated by varying early innate immune responses. Therefore, we started dissecting immune mechanisms that mediate the anti-fentanyl vaccine antibody responses. We characterized the effect of fentanyl vaccine formulated with INI-4001/alum, or INI-4001 liposomes on the activation, maturation, and phenotypic differentiation of two major antigen presenting cells subsets (APCs) using *in vitro* culture of murine JAWS II DCs (Figure 9) and RAW 264.7 macrophage cells (Figure 10). All formulations enhanced DCs surface expression of CD86 and MHC II compared to the vehicle control. In line with the *in vivo* testing in mice and rats, F-CRM adjuvanted with INI-4001/alum and cationic INI-4001/DOPC/DC-Chol liposomes significantly and equally enhanced the expression of the co-stimulatory molecule CD86 (Figure 9A) and the maturation marker MHC II (Figure 9B) over neutral, and anionic liposomes. The cationic liposomal adjuvanted vaccines were the most prominent in triggering the expression of the activation marker CD40 compared to all other formulations, followed by the INI-4001/alum adjuvanted vaccine. Notably, both neutral INI-4001/DOPC/Chol and anionic INI-4001/DOPG/Chol liposomes exhibited only slight to no impact on the expression of CD40.

**Figure 9.**
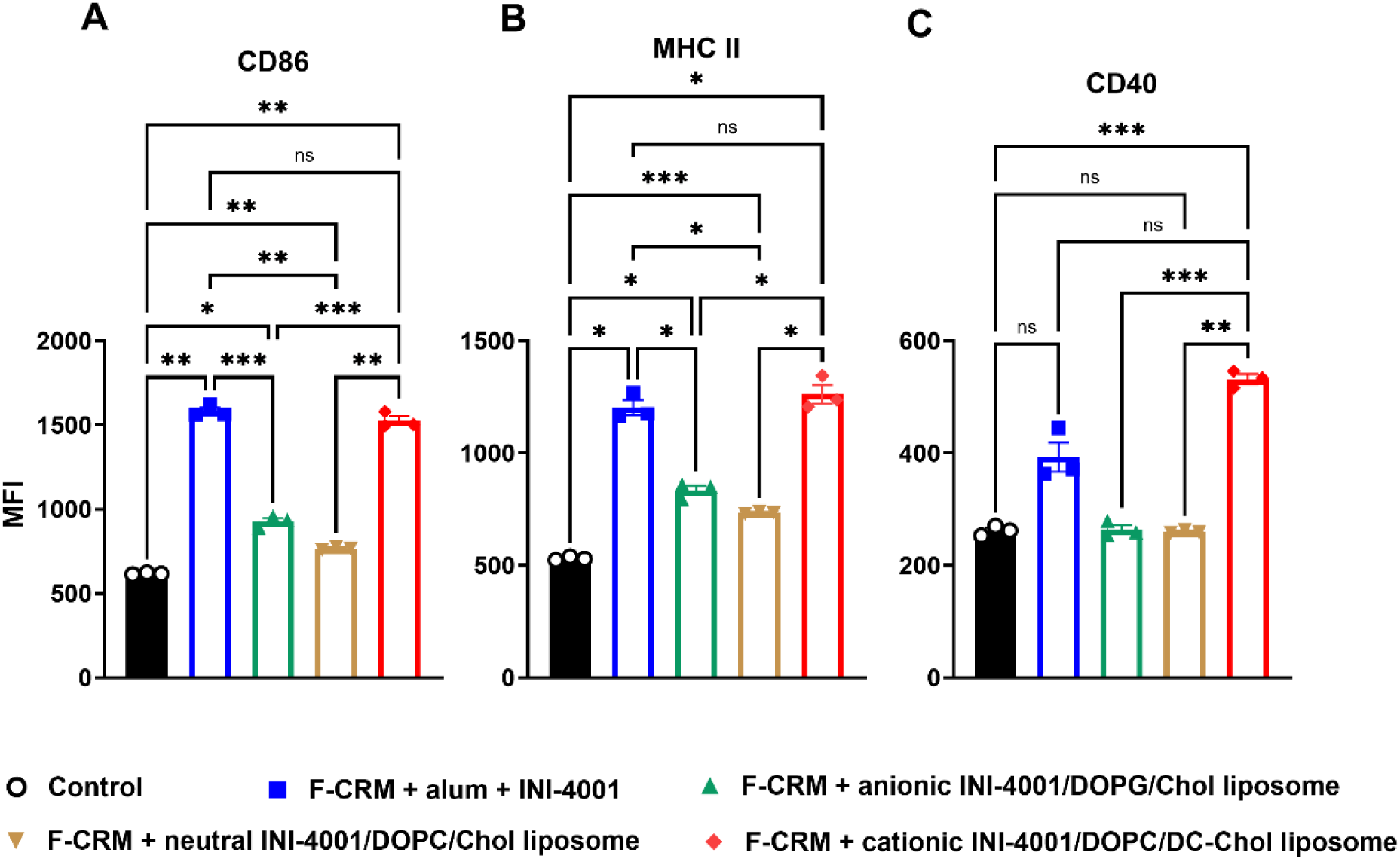
Effects of anti-fentanyl vaccine formulations on *in vitro* activation of DCs. JAWS II cells were primed either with F-CRM and INI-4001/alum, F-CRM + INI-4001 liposomes, or media as a control. Twenty-four hours later, cells were harvested, stained and processed for flow cytometry analysis. Cells were assessed for the effect of vaccines on inducing DC co-stimulatory molecules and maturation markers (A) CD86, (B) MHC II and (C) CD40. MFI: median fluorescence intensity. Data are representative of plots from one of two independent experiments. The error bars are standard deviation. Statistical analysis was performed using Brown-Forsythe and Welch’s ANOVA test. **p < 0.001, ***p < 0.0005, ****p < 0.0001.

**Figure 10.**
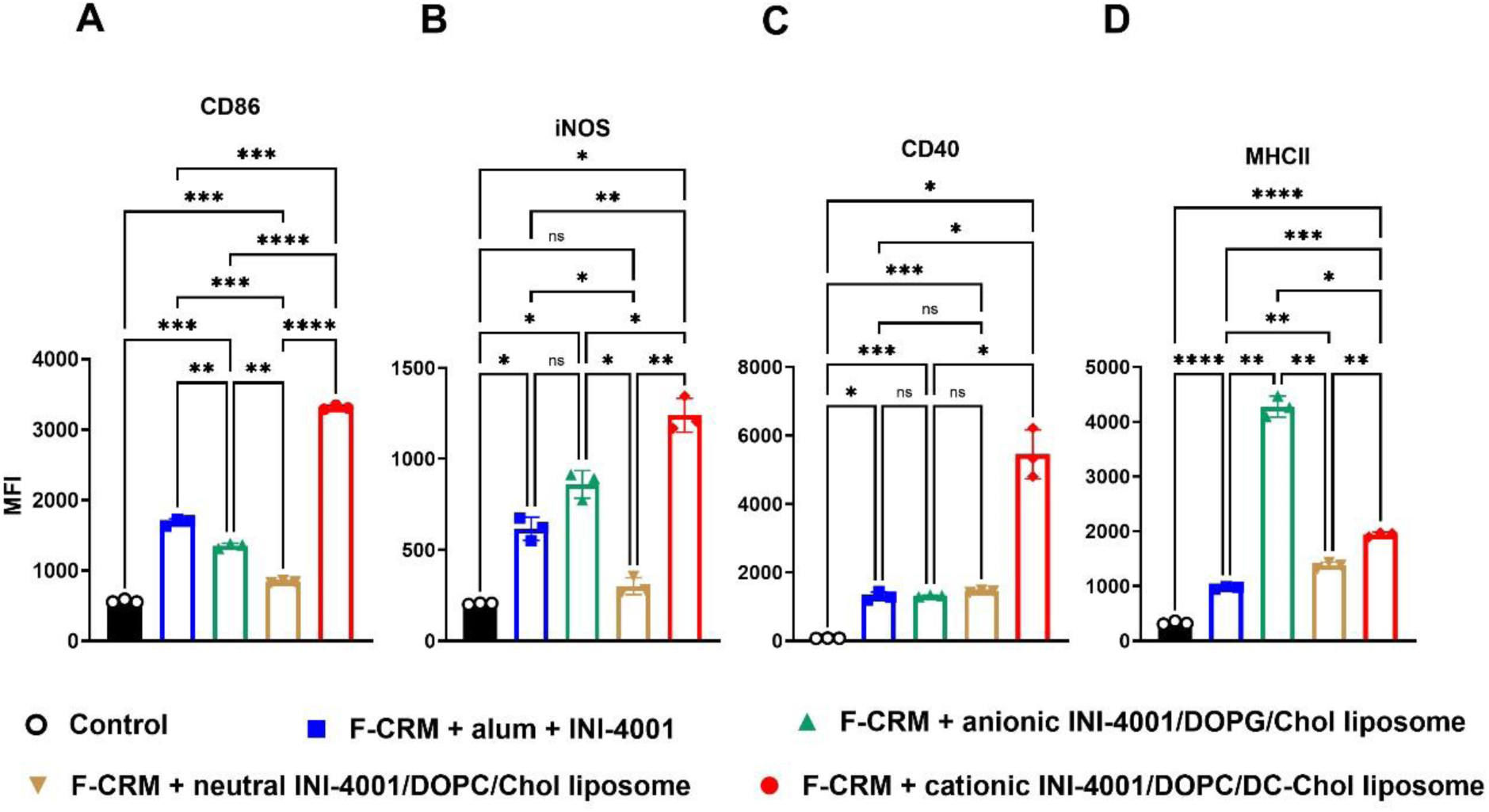
Effects of anti-fentanyl vaccine formulations on *in vitro* activation of macrophages. RAW264.7 cells were primed with F-CRM and INI-4001/alum, F-CRM + INI-4001 liposomes, or media as a control. Twenty-four 24 hours later, cells were harvested, stained, and processed for flow cytometry analysis. Cells were assessed for the effect of vaccines on inducing the expression of macrophages co-stimulatory molecules, activation and maturation markers (A) CD86, (B) inducible nitric oxide synthase (iNOS), (C) CD40 and (D) MHC II. Median fluorescence intensity (MFI). Data are representative of plots from one of two independent experiments. The error bars are standard deviation. Statistical analysis was performed using Brown-Forsythe and Welch’s ANOVA test. **p < 0.001, ***p < 0.0005, ****p < 0.0001.

In addition, we explored the effect of these formulations on macrophages, one of the major innate immune cell populations affecting downstream vaccination outcomes, using murine RAW264.7 cell line (Figure 10). Consistent with DC profiling, macrophages stimulated by cationic INI-4001/DOPC/DC-Chol liposomal adjuvanted vaccine expressed the highest levels of the co-stimulatory markers CD86 and CD40 and the activation marker iNOS compared to all other formulations. F-CRM/alum/INI-4001 ranked second in terms of CD86 expression and was equivalent to anionic INI-4001/DOPG/Chol liposomes with respect to iNOS upregulation. INI-4001/alum, anionic, and neutral liposomes induced comparable expression levels of CD40. All INI-4001 liposomal formulations induced significantly higher levels of the maturation marker MHC II compared to the INI-4001/alum combination, with the anionic liposomes being superior. Anionic INI-4001/DOPG/Chol liposomes and neutral INI-4001/DOPC/Chol liposomes were equally ineffective when tested on DCs. In contrast, the anionic INI-4001/DOPG/Chol liposomes were more effective in inducing phenotypic differentiation of macrophages. This could be attributed to variations in the uptake efficiency of particles with opposite surface charges in different cell types. Despite the statistically significant increase in the expression of activation and maturation markers in RAW264.7 cells treated with double adjuvanted vaccine (INI-4001/alum) compared to media only group, this boost was mild compared to the effect of positively charged liposomes in macrophages. These data suggest that DC activation provides a better biomarker predictive of vaccine performance *in vivo*. These data also imply that different adjuvants have varying effects on different types of cell subsets. These *in vitro* assays provide cellular mechanistic insights into adjuvants and vaccines and can be utilized for early screening of effective vaccine formulation during early-stage development. Overall, these studies urge us to further investigate the *in vivo* innate immune response associated with these liposomes to better understand the innate immune cell subsets and their contributions to vaccine efficacy.

## 4. Discussion

The holy grail of anti-drug vaccine efficacy is the quantity and the quality of antibodies response^33^. Considering the high lipophilicity and potency of fentanyl, high fentanyl-specific antibodies titer with high binding avidity (low Kd) is a prerequisite to block fentanyl entry to the brain. Such a favorable vaccination outcome requires a robust initial innate immune response to trigger an optimal CD4^+^ T cell-dependent B cell-specific immunity^34^. Upon exposure to danger signals, i.e., TLR7/8 agonists, DCs undergo a maturation process characterized by the increased formation of MHC-peptide complexes and the up-regulation of costimulatory molecules (CD86, CD40) as well as cytokines secretion that drives Th1 response^35^.

In this study, we explored the effect of liposomal formulations of INI-4001 and their surface charge on immunogenicity and efficacy of anti-fentanyl vaccines, a promising vaccine candidate to counteract deliberate and accidental fentanyl exposure-related overdose and OUD^6, 25^. INI-4001 is a novel lipidated oxoadenine TLR7/8 agonist that was previously shown to significantly enhance the immunogenicity and efficacy of the anti-fentanyl vaccine when adsorbed on Alum^11, 12^. Of note, the phospholipid attached to INI-4001 enables direct adsorption of this adjuvant to alum. In addition, the phospholipid was designed to allow for the incorporation of INI-4001 into liposomes with the TLR7/8 signaling segment displayed on the surface of these spherical lipid bilayer vesicles. Liposomes represent a well-established delivery system for antigens and adjuvants due to their unique properties, such as high versatility, biocompatibility, and safety profile^36–38^. Previously, liposomal formulations of small molecule adjuvants like TLR7/8 agonist (3M-053) and TLR4 agonist (MPLA) have been shown to potentiate their immune responses, limiting their systemic distribution and side effects^39–41^. Owing to the high flexibility of liposomes, their electrical surface charge can be modulated by incorporating cholesterol derivatives and phospholipids with different charges to produce negative (anionic), neutral, or positive (cationic) liposomes^42^. Modulating the physico-chemical properties of liposomes, e.g., size and surface charge, can greatly influence their immunostimulatory activity, stability, and antigen loading capacity^37, 43, 44^. Here, all liposomes were made at the same particle size range of 50-60 nm in diameter to study the impact of liposome surface charges, which is considered a major factor determining cellular uptake, protein binding, antigen adsorption, and depot effect^36–38^.

The interaction between hapten-protein conjugates and the liposomal adjuvants is crucial for vaccine efficacy as studies showed the co-delivery of antigen and adjuvant plays an important role in eliciting a robust immune response^32, 38, 45^. Therefore, characterizing the extent of protein conjugate (antigen) adsorption to the delivery system is essential. CRM_197_ protein has an isoelectric point of ∼5.7 and is negatively charged at a physiological pH of 7.4, this surface charge will dictate its adsorption to various liposome formulations. This is evidenced in the case aluminum salts-based adjuvants, where positively charge alum is well suited for adsorption of negatively charged antigens, while Adju-Phos^®^ has a net negative charge at neutral pH, effectively adsorbs positively charged antigen^46^. Similarly, in our study, we expected negatively charged F-CRM to show greater adsorption to positively charged INI-4001/DOPC/DC-Chol liposomes compared to neutral and negatively charged ones.

Using the analytical ultracentrifugation technique, we initially observed identical sedimentation patterns across a 3-fold concentration range for all liposome formulations. This suggests that the liposomes exhibit stable behavior in solution within this concentration range, with negligible mass action effects. The incorporated fluorescently labeled F-CRM in cationic and neutral liposomes tends to sediment slower than the UV-measured empty liposome controls. To explain this observation, it is important to realize that UV measurements do not reflect absorbance proportional to mass, but rather light scattering proportional to the sixth power of the radius of the particles. As a result, larger particles are emphasized, while smaller ones are underrepresented. In contrast, fluorescence signals are directly proportional to the number of fluorophores, capturing all labeled F-CRM molecules, including those incorporated in smaller liposomes. Consequently, the reduced sedimentation coefficient (s-value) distribution for the FITC F-CRM:INI-4001 liposome mixtures, compared to the UV-measured empty liposome controls, is attributed to the inclusion of smaller liposomes in the fluorescence-based measurements.

Characterization of FITC labeled F-CRM using fluorescence optics confirmed that the fluorescent labeling did not significantly alter the hydrodynamic properties of F-CRM. The protein remained monomeric at 1.6 µM, consistent with its predicted molecular mass based on its amino acid sequence. Analysis of the raw fluorescence data comparing FITC labeled F-CRM alone and in combination with liposomes suggests fluorescence quenching occurs as a function of liposome charge. Since all samples were measured under identical conditions—same photomultiplier tube gain, voltage settings, and fluorescent material concentration (1.6 µM FITC F-CRM)—the observed differences in fluorescence intensity are attributable to quenching. The FITC labeled F-CRM control, in the absence of liposomes, exhibited the highest fluorescence yield, while the cationic liposomes showed the lowest fluorescence signal, indicating the strongest quenching effect. Quenching is proportional to the integration efficiency of FITC F-CRM and could be a result of the altered chemical environment of the fluorophore when bound by the cationic lipids. Furthermore, the sedimentation profiles of the fluorescently labeled liposomes indicated varying degrees of FITC F-CRM incorporation based on liposome charge. Cationic liposomes demonstrated the highest level of F-CRM binding, followed by neutral liposomes, while anionic liposomes showed no detectable binding of the protein. Together with the fact that F-CRM/INI-4001 are entirely bound to alum suggests that both delivery platforms (alum and cationic INI-4001/DOPC/DC-Chol liposomes) through adsorption can effectively co-deliver the antigen and adjuvants to APCs, an attribute that demonstrated to be crucial for effective immune response^32, 38, 47^.

In the field of cancer, positively charged liposomes have been shown to enhance DCs activation, as many studies on mice and *in vitro* human DCs demonstrated elevation in maturation markers and proinflammatory cytokines profiles by cationic liposomes. In contrast, anionic liposomes in mice DC *in vitro* culture were shown to promote DC resistance to maturation and inefficient activation of T cell responses^48–50^.

Cationic liposomes, due to their positively charged nature, are readily attracted to the anionic membrane of APCs, facilitating the uptake and delivery of their antigens and adjuvants cargo via the endosomal route, leading to antigen presentation on MHC class II molecules. Thus, cationic liposomes can help initiate and promote adaptive antigen-specific antibody responses^51, 52^. Moreover, cationic liposomes were shown to form a depot at the injection site regardless of the size, perhaps due to interaction with negatively charged interstitial proteins^53^. In agreement with this, our *in vivo* data in mice and rats showed that INI-4001 cationic liposomes significantly enhanced the immunogenicity and efficacy of the anti-fentanyl vaccine compared to the anionic and neutral liposomes. Previously we demonstrated that the efficacy of alum adjuvanted F-CRM vaccine that is primarily IgG1 mediated, can be enhanced by depleting IL-4 or incorporating TLR7/8 agonist, skewing the IgG pool toward a more balanced IgG1/IgG2a response^20, 21, 54^. Similarly, here cationic liposomal formulations of INI-4001 enhanced the immunogenicity of F-CRM in mice and induced a significant shift in IgG1/IgG2a antibody titer ratio to a more biased Th1 response. In both cases, the enhancement in efficacy observed was associated with increased total IgG titer, and increased class switching from IgG1 to IgG2a, creating a balanced Th1/Th2 response in the former case or Th1 biased CD4^+^ T cells responses in the latter. These findings further support our previous reports that specific IgG subclasses are irrelevant for vaccine efficacy and that antibody-mediated effector mechanisms do not play a role in opioid specific antibody function^55, 56^. We conducted further tests to evaluate the efficacy of INI-4001 liposome formulations in rats. *In vivo* rat studies showed that both F-CRM/alum/INI-4001 and F-CRM + cationic INI-4001/DOPC/DC-Chol liposomes significantly induced serum fentanyl-specific IgG antibody titer, and equally blocked the distribution of 0.3 mg/kg fentanyl to the brain. This observed superior immunogenicity and *in vivo* efficacy of F-CRM + cationic INI-4001/DOPC/DC-Chol liposomes and F-CRM/alum/INI-4001 is proposed to be due to their ability to co-deliver antigen and adjuvants to APCs and therefore significantly activate DCs and macrophages resulting in fully mature innate immune cells that can effectively initiate and sustain robust adaptive antibody responses.

Overall, vaccine adjuvant platforms that can enhance targeted delivery, minimize systemic distribution and reactogenicity, and enable efficient antigen adsorption and modulation of immune response are of high interest. Incorporating a TLR7/8 ligand into cationic DOPC/DC-Chol liposomes presents a potent adjuvant system that could aid in developing effective vaccines against OUD and possibly other SUD.

## Abbreviations

(TLR): Toll-like receptor
(F): fentanyl hapten
(APC): antigen-presenting cells
(DC): dendritic cells
(OUD): opioids use disorders
(IL-4): Interleukin 4
(MORs): μ-opioid receptors

## ACKNOWLEDGEMENTS

This work was supported by HHSN272201800048C (J.E.), 75N93020C00039 (J.E.), UG3DA048386 (M.P.), F31 DA059235 (F.A.H.), C150-2017-00015 (B.D.), 1R01GM120600 (B.D.) and DG-RGPIN-2019-05637 (B.D.). The Canadian Center for Hydrodynamics is funded by the Canada Foundation for Innovation grant CFI-37589 (B.D.). Research reported in this publication is solely the responsibility of the authors and does not necessarily represent the official views of the National Institutes of Health.

## AUTHOR CONTRIBUTIONS

F.A.H., N-M.N.L., D.S., H.A., L.H., S.B., K.S., B.D., J.T.E., D.B., and M.P. conceived and designed the work. F.A.H., N-M.N.L., D.S., H.A., L.H., S.B., K.S., B.H. acquired and analyzed data. All authors contributed to data interpretation. F.A.H. and N-M.N.L. drafted the manuscript. D.B., J.T.E., and M.P. provided substantive revisions. All authors reviewed, revised, and approved the submitted manuscript.

## Conflict of Interest

M.P. is the co-inventor of patents disclosing fentanyl haptens reported in this manuscript (F1 or F) fentanyl hapten conjugates (F1-CRM, F-CRM), and methods for using them. D.B. and J.T.E. are co-founders, employees, and shareholders of Inimmune Corporation, which owns an exclusive license for INI-4001. M.P., D.B. and J.T.E. are co-inventors of patents disclosing formulations of fentanyl vaccines formulated with INI-4001. M.P. is the founder and shareholder of CounterX Therapeutics. All other authors declare no competing interests.

