## Supplemental materials for "A cationic liposome-formulated Toll Like Receptor (TLR)7/8 agonist enhances the efficacy of a vaccine against fentanyl toxicity"

**Supplementary information**

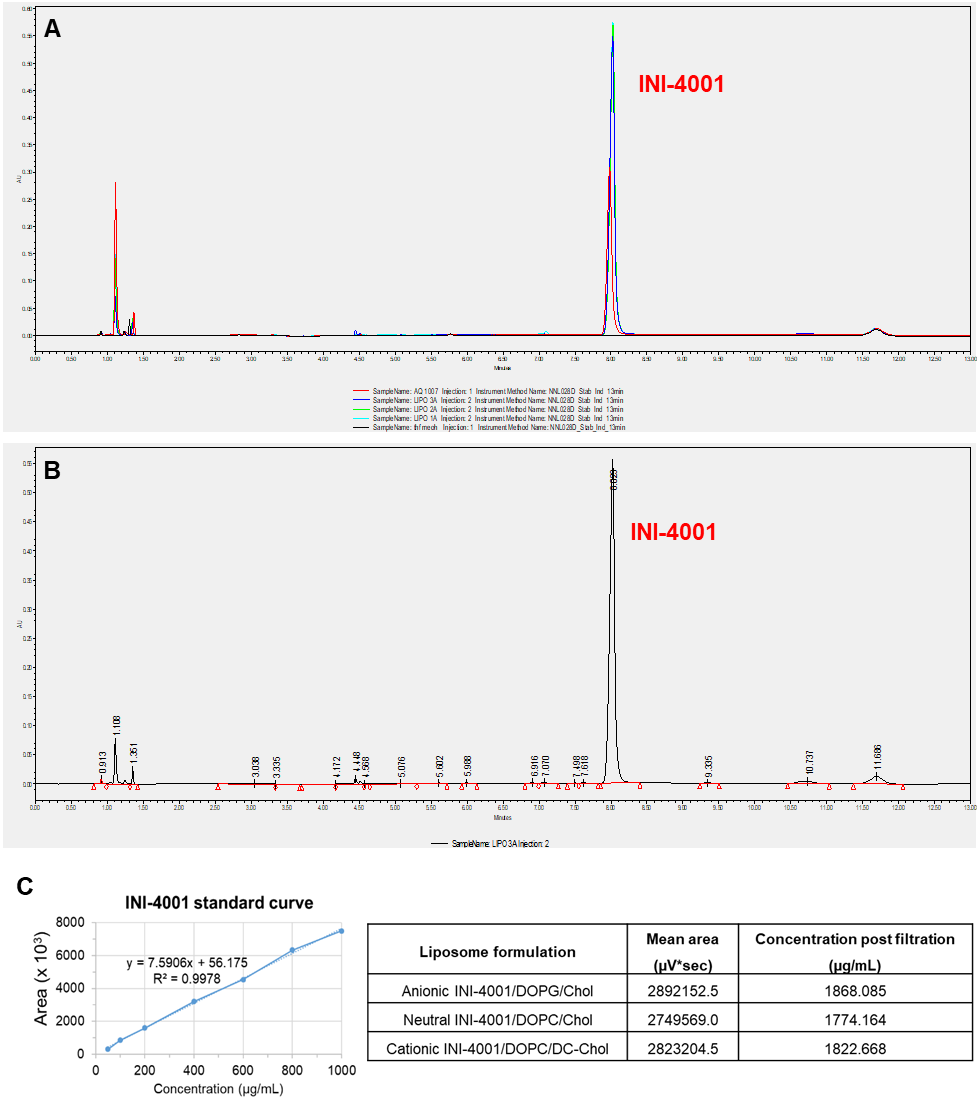

**Figure S.1. UHPLC analysis of INI-4001 in liposomes.** (A) Chromatograph showing the distinct peak of INI-4001 in differently charged INI-4001 liposomes and aqueous INI-4001 formulations. (B) Typical integration of the chromatographic peaks in a liposome formulation. The retention time of INI-4001 was ~8.02 minutes. (C) Linearity assessment of INI-4001, established within the range of 50-800 µg/mL (left) and the quantitative concentration of INI-4001 in liposomes post filtration (right).

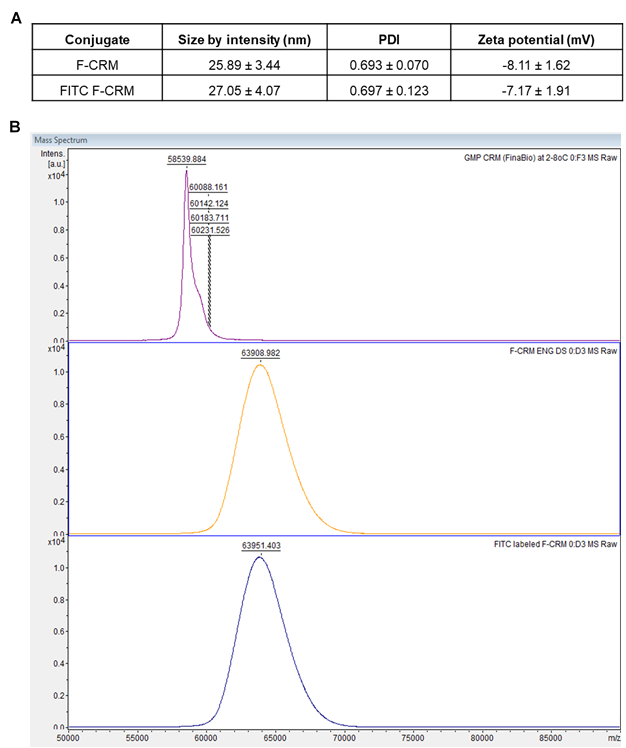

**Figure S.2. Characterization of FITC labeled F-CRM compared to F-CRM.** (A) Size, PDI, and zeta potential assessment by DLS. (B) Molecular weight assessment by MALDI-TOF.

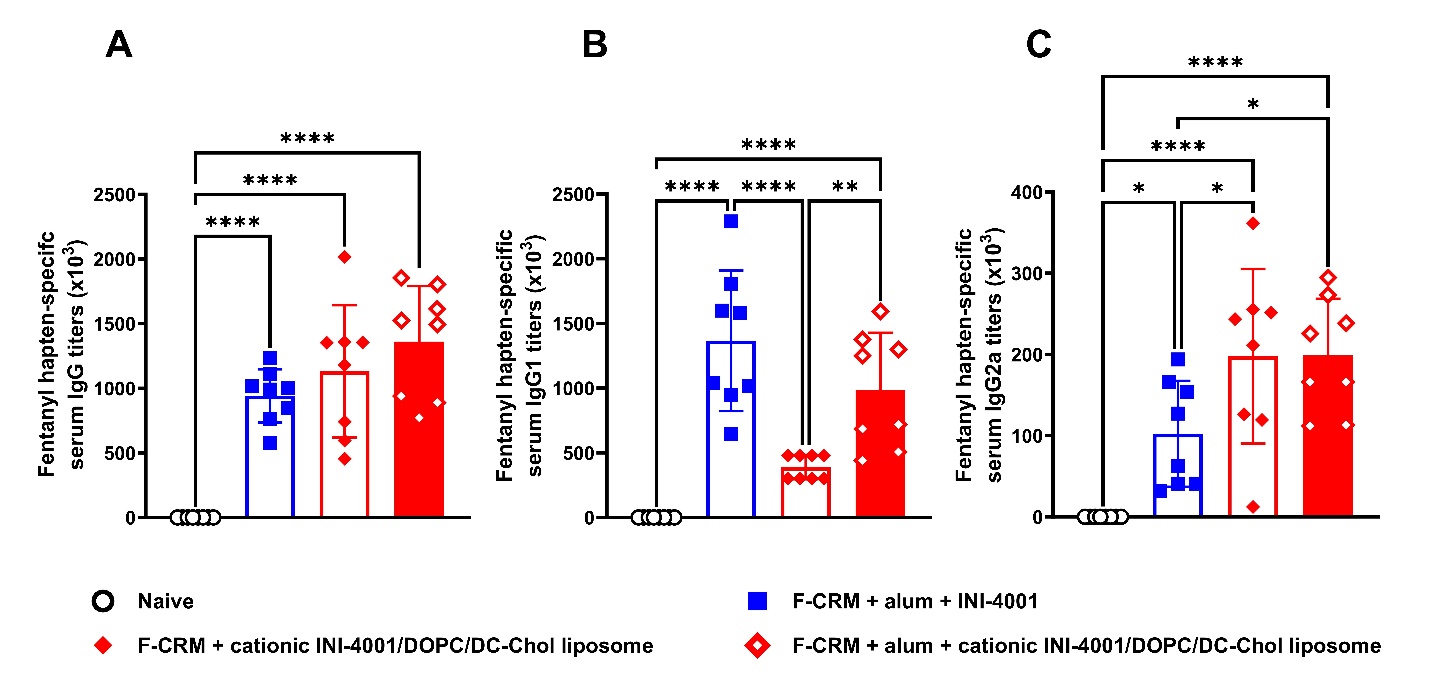

**Figure S.3. The immunogenicity and Ig subclass when adjuvanting F-CRM with cationic INI-4001/DOPC/DC-Chol liposomes with and without alum in comparison with aqueous 4001 + alum formulation in mice.** Six- to eight-week-old female BALB/c mice were vaccinated on days 0 and 14, IM, with 5 µg F-CRM plus 10 µg INI-4001 and 22.5 µg alum. The serum was collected on day 21. Anti-fentanyl hapten IgG (A), IgG1 (B), and IgG2a (C) antibody concentrations were measured via ELISA. Statistical analysis was conducted using one-way ANOVA and Tukey’s multiple comparisons. *p ≤ 0.05, **p ≤ 0.01, ***p ≤ 0.001, ****p ≤ 0.0001; color of asterisks indicates comparison group.

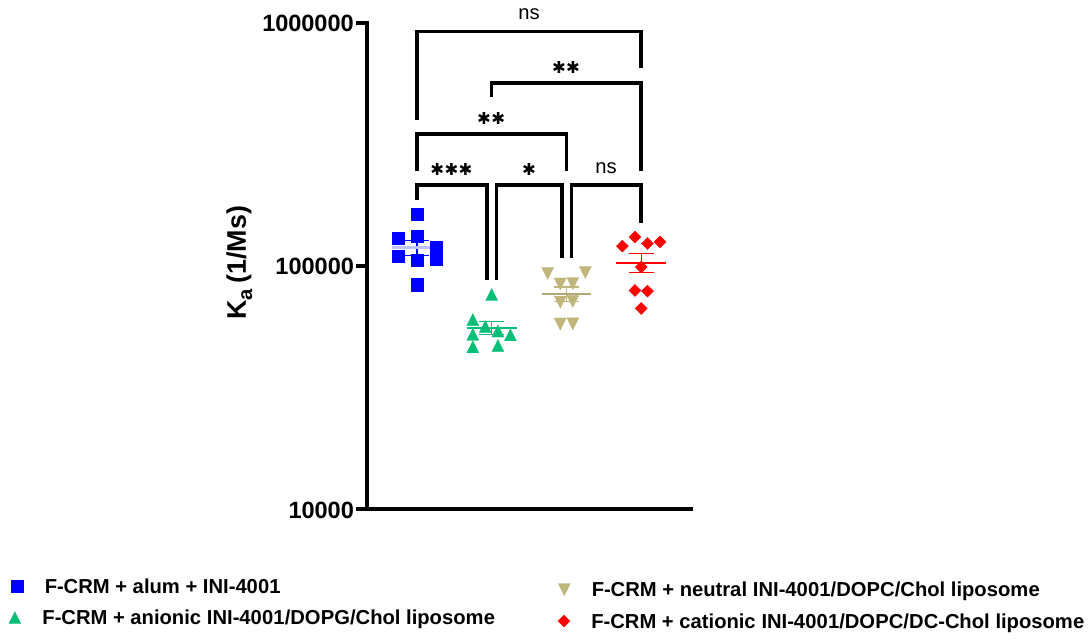

**Figure S.4.** **The antibody association rate constant (*K*_ass_) to fentanyl hapten**. Mouse serum samples were used to measure the apparent association rate (*K*_ass_) between F_1_ and polyclonal mouse serum for each individual serum sample using Octet SA Biosensors (Sartorius 18-5019). Association rate constants (*K*_ass_) were calculated by processing raw data using Sartorius Analysis Studio v13.0.1.35. K_a_ values from curves with R^2^>0.95 were plotted in GraphPad Prism v10.3.1.  Statistical significance was determined using a one-way ANOVA with Dunnett's multiple comparisons test. *p ≤ 0.05, **p ≤ 0.01, ***p ≤ 0.001; color of asterisks indicates comparison group.
